# Minimally invasive serial collection of cerebrospinal fluid reveals sex-dependent differences in neuroinflammation in a rat model of mild traumatic brain injury

**DOI:** 10.1101/2024.02.26.582018

**Authors:** Josh Karam, Nimrah Ashfaq, Cynthia Benitez, Victor Morales, Elizabeth Partida, Michelle Hernandez, Jordan Yokoyama, Alyssa Villegas, Brielle Brown, Pooja Sakthivel, Aileen J. Anderson, Brian J. Cummings

## Abstract

Traumatic brain injuries (TBI) are the seventh leading cause of disability globally with 48.99 million prevalent cases and 7.08 million years lived with diability. Approximately 80% of TBI patients are diagnosed with mild TBI (mTBI), or concussion, caused by nonpenetrating mechanical trauma to the head or body along with sudden rotational motion of the head. Studies investigating the temporal dynamics of neuroinflammation after mTBI are greatly needed. Without longitudinal studies, translating preclinical studies to clinical studies remains challenging as the difference in timing remains poorly understood. In this study, we describe a method of minimally invasive serial cerebrospinal fluid (CSF) collection that enables longitudinal investigation of CSF inflammation. The method described in this study can easily be adapted by any laboratory prepared for animal studies. Multiplex immunoassay of serially collected and singly collected CSF samples show collection frequency does not alter protein expression in the CSF. Further, sex-dependent differences in TBI have been reported, but remain poorly understood. This study establishes a framework for assessing sex difference in neuroinflammation after a concussion. We showed that results vary based on the framing of the statistical test. However, it is evident that males experience a more robust inflammatory response to a single concussion than females.

## 1. Introduction

Traumatic brain injuries (TBI) are the seventh leading cause of disability globally with 48.99 million prevalent cases and 7.08 million years lived with diability^1^. Approximately 80% of TBI patients are diagnosed with mild TBI (mTBI), or concussion, caused by nonpenetrating mechanical trauma to the head or body along with sudden rotational motion of the head^2–4^. mTBIs commonly occur in falls, motor vehicle crashes, contact sports, military duty, and incidences of domestic violence^2,5–7^. While the typical prognosis of a single concussion is positive, with full recovery within 3 months^3^, approximately 15 percent of patients also experience post-concussion symptoms (PCS) that are exacerbated with repeated mTBIs^3,6,8,9^

The pathophysiology of mTBI remains poorly understood, primarily due to heterogeneity in presentation and outcome. mTBI is associated with structural abnormalities, excitotoxicity, metabolic stress, and increased inflammation^10–12^. However, much of our understanding of mTBI pathophysiology stems from more severe injuries. Major distinctions between mTBI and more severe TBI include the absence of intracranial lesions, hemorrhages, and skull fractures in mTBI^5,6,8,13^. Further, there are differences in the inflammatory response based on injury severity. In a study analyzing whole blood from pediatric TBI patients, severe TBI caused greater concentrations of TNFα, IL-6, IL-8, and IL-10 compared to mTBI. However, mTBI patients experience a greater concentration of IFNγ and IL-17A^14^. While some overlap of the neuroinflammatory response across TBI severities is expected, studies investigating the temporal dynamics of neuroinflammation after mTBI are still needed to uncover the mechanisms behind PCS, as well as to identify potential biomarkers that can be leveraged to develop more reliable point-of-care diagnostics and therapeutic strategies. Furthermore, identified mechanisms and biomarkers will need validation in both sexes.

There have been many reported sex-dependent differences in mTBI^5,6,9,15,16^. Interrogating sex differences in mTBI is important simply because TBI outcomes exhibit significant sex differences with women experiencing lower incidences of morbidity and mortality after TBI than age-matched men^17^. In a recent publication, Liu *et al.* reported that males, but not females, treated with minocycline after a repeat mTBI (rmTBI) experienced an increased cold thermal threshold and decreased anxiety-like behavior. There was also sex differences in the microglial response to rmTBI as both males and females exhibited increased microglia and brain-derived neurotrophic factor (BDNF) mRNA in the anterior cingular cortex, but only males saw an increase in BDNF in the nucleus accumbens^18^. Another sex difference noted in the literature is the effect of regulatory T cells (Tregs) on microglial activation. In a hypoxia-ischemia model of brain injury, Beckmann *et al.* found increased numbers of Tregs and decreased microglial activation in female brains, but in male brains they found Tregs promoted vascular injury^19^. While this study was not in TBI, it highlights the sex-differences in the innate and adaptive immune responses, which are hypothesized to be affected by sex hormones. Estrogen and progesterone have been shown to be neuroprotective after TBI, reducing mortality and improving neurological outcomes, by reducing neuronal apoptosis, inhibiting microglial and astrocytic activation, and reducing brain edema^17,20–22^. To fully understand the inflammatory response to TBI, it is vital to assay cerebrospinal fluid (CSF), as it has been shown that CSF biomarkers such as MAP-2 enhance prognosis capabilities^23^. The literature analyzing the CSF proteome over time after TBI is very limited, primarily due to the lack of a CSF collection method that is minimally invasive and enables serial collections^24–27^.

This study has three primary goals: 1) establish a method of minimally invasive serial CSF collection that can be easily utilized by any laboratory equipped for animal studies without any specialized equipment, 2) determine if serial CSF collection alters CSF protein detection, and 3) provide a framework for interrogation of sex differences in CSF proteomics after neurotrauma.

## 2. Methods

### 2.1. Animal care and concussion

All experiments, including animal housing conditions, surgical procedures, and postoperative care, were in accordance with the Institutional Animal Care and Use Committee guidelines at the University of California–Irvine (UCI). Eighteen male and eighteen female Long-Evans rats (200-225g, Charles River Laboratories, San Diego, CA) were randomly divided into six experimental groups: female concussion with single CSF withdrawal, female concussion with serial CSF withdrawal, female sham with serial CSF withdrawal. male concussion with single CSF withdrawal, male concussion with serial CSF withdrawal, and male sham with serial CSF withdrawal. Each group had n=6 rats.

A pneumatic controlled cortical impact device (TBI-0310 Head Impactor, Precision Systems and Instrumentation, LLC, Fairfax Station, VA) was used for concussion. A foam bed modified from “Marmarou foam” (Type E Bed, Foam to Size, Inc., Ashland, VA) was modified to fit on the stage of the CCI device, and a trench was cut out to position tubing and a nose cone to keep the animal under anesthesia for the duration of the procedure. All animals spent 10 min under anesthesia, including shams; 5 min induction at 3.5% isoflurane, and 2 min to mark the tail, shave the head, and apply ophthalmic ointment. The animal was placed onto the modified Marmarou foam and positioned to breathe 3.5% isoflurane from a nose cone. Lab tape was lightly applied to the ears and fixed to the foam to position the head level and prevent movement during breathing. The rat was at the CCI station for 3 min until a 5-mm diameter probe tip was used to deliver an impact with speed 5.0 m/s, 1.0 mm depth, and 50 ms dwell time. In total, rats are under anesthesia for 10 min from knockdown to TBI impact. The isoflurane percentages were designed to minimize head movement from respiration while zeroing the piston and during impact while still keeping the animal unresponsive to a toe pinch reflex. Following impact, animals were moved to a recovery area where righting time was recorded. Sham animals underwent the same 10-min procedure with anesthesia but were not impacted.

### 2.2. Cerebrospinal fluid sample collection and storage

This protocol is adapted from Han *et al.* making the procedure easy to adapt for any laboratory without the use of custom built equipment or specialized tools^27^. To prep for CSF collection, an isofluorane nose cone was taped to the short side of a rectangular brick (can be any raised platform such as a pipette tip box) so the rat’s head could be positioned vertically while breathing in isofluorane.

Before CSF collection, Long-Evans rats were anesthetized using an isofluorane induction chamber with 3.5% isofluorane and shaved. For serial collections, the shaved area was sterilized with ethanol and betadine. After the rat is placed on the brick with the nose in the nose cone, the rat’s head was positioned vertically. The rat skin is then pulled back to ensure the area around the foramen magnum is tight, and tape is placed over the shoulder blades.

To start the CSF collection, a sterile hemostat was used to bend the needle of the 27G winged butterfly needle approximately 90 degrees so the bent part is about 4mm (Fig 2A). Before insertion of the needle, the foramen magnum needs to be located. The external occipital protuberance was used as a landmark. This should feel like a ridge on the skull. About 6mm past the ridge is the foramen magnum (Fig 2B). The bent 27G winged needle was inserted into the foramen magnum (an extremely soft spot) with no resistance. Once inserted, the syringe was pulled to 0.1 mL, and clear CSF fluid flowed into the tubing of the needle (Fig 2C). Once CSF stopped flowing, the needle was removed. There were no visible signs of damage besides the insertion point (Fig 2D). The CSF was then transferred from the tubing into a microcentrifuge tube on ice (Fig 2E). CSF samples were measured using a pipette then supplemented with 1X protease/phosphatase inhibitor cocktail (Cell Signaling Technologies, #5872). CSF was stored at −80℃.

### 2.3. Luminex xMAP multiplex immunoassay

CSF Samples were thawed on ice and centrifuged. A bead-based multiplex cytokine kit (LXSARM-17, R&D Systems, Minneapolis, MN, USA) was used to measure CSF levels of CXCL2, CXCL3, ICAM-1, IFN-γ, IL-1α, IL-1β, IL-2, IL-4, IL-6, IL-10, IL-13, IL-18, L-Selectin, TIMP1, TNF-α, and VEGF. This kit was chosen due to its published use in rat TBI studies^28,29^. All samples were run in duplicate. Results were analyzed using mean fluorescent intensities as previously reported^30,31^.

### 2.4. Immunohistochemical labeling of Iba1 and GFAP

Brain tissue sections were immunohistochemically labeled with primary antibodies for Iba1 (1:500, 019-19741, FujiFilm Wako) and GFAP (1:1500, Z033429-2, Agilent). Biotinylated donkey anti-rabbit (1:500, 715-066-152, Jackson ImmunoResearch) secondary antibodies were used, followed by VECTASTAIN ® Elite® ABC Kit Peroxidase (PK-6100, Vector Laboratories) and DAB Substrate Kit, Peroxidase with Nickel (SK-4100, Vector Laboratories) according to manufacturer’s instructions.

### 2.5. Stereological quantification of microglia and astrocytes

Stereological quantification was performed on every 12th brain tissue section with thickness of 30um. The corpus collosum was traced at the connection between the left and right cerebral hemisphere. An average of 10 counting frames measuring 80 µm x 80 µm was implemented. The systematic random sampling (SRS) grid measured 250 µm x 250 µm. The optical dissector height consisted of a guard zone thickness of 1um and a dissector height of 18um. The CE (Coefficient Error) was 0.1.

### 2.6. Semi-quantitative assessment of microbleed presence

A categorical scale of 0-3 was implemented to rate the presence of microbleeds at three regions of the brain: cerebral cortex, corpus collosum, and hippocampus. A score of 0 indicates no visible hemosiderin granules. A score of 1 indicates that a few hemosiderin granules were visible. A score of 2 indicates that some hemosiderin granules were visible. A score of 3 indicates that many hemosiderin granules were present. The scores for each region were summed up for each section. 5 serial sections were used for each animal. The total sum of all 5 sections was used for statistical analysis.

### 2.7. Statistical analysis

For analysis of serially collected samples, a two-way mixed effects analysis with a post-hoc Holm-Šídák’s multiple comparisons test was used. For statistical tests that did not require matching samples, an ordinary two-way ANOVA with a post-hoc Holm-Šídák’s multiple comparisons test was used.

## 3. Results

### 3.1. CSF was collected serially from rats with no evidence of negative side effects

CSF was collected from a total of 36 rats (18 male and 18 female). 24 rats underwent serial collections 1 day before concussion and then 1 day post injury (dpi), 3 dpi, 7 dpi, and 14 dpi. The remaining 12 rats underwent a single CSF collection at 7 dpi only (Figure 1). No rats exhibited signs of pain/distress, weight loss, infection, or swelling. There was no attrition of animals due to mortality. If CSF samples were contaminated with blood upon collection, they were immediately centrifuged for 30 seconds to separate the CSF from blood, yielding clear CSF samples. Centrifugation has previously been shown not to affect CSF composition^32^. Of the 132 attempts to collect CSF samples, 121 attempts (92%) resulted in clear CSF (Figure 2 A-E).

**Figure 1:**
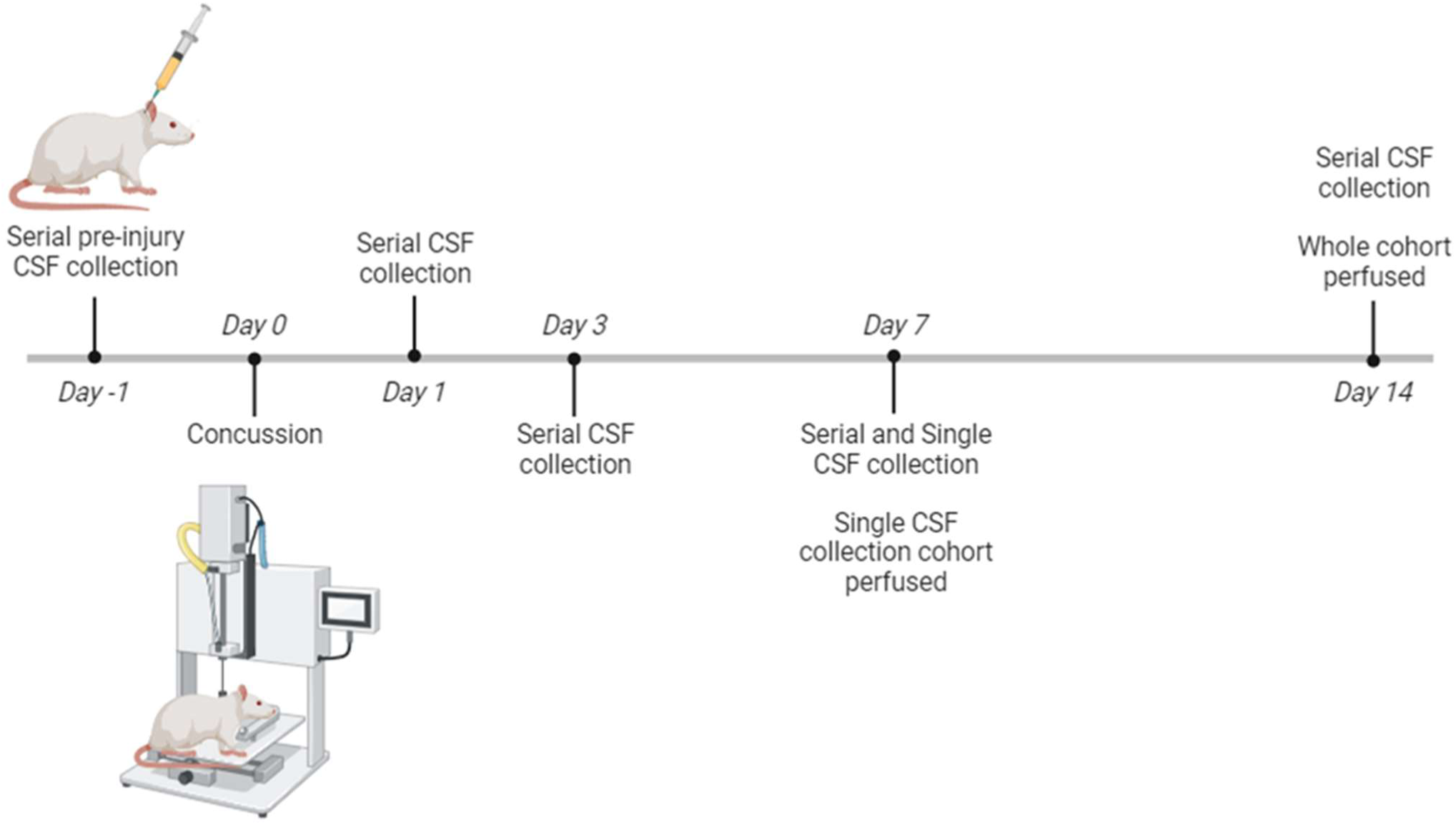
Experimental timeline schematic. Briefly, pre-injury CSF samples were collected 1 day before injury, except from the single collection CSF group. On day 0, all groups of rats were given either given a concussion or were shams. Serial collections of CSF were taken at 1 day, 3 days, 7 days, and 14 days post­injury. At 7 dpi, a group of rats with injury underwent a single CSF collection and were sacrificed. All other groups were sacrificed at 14 dpi.

**Figure 2:**
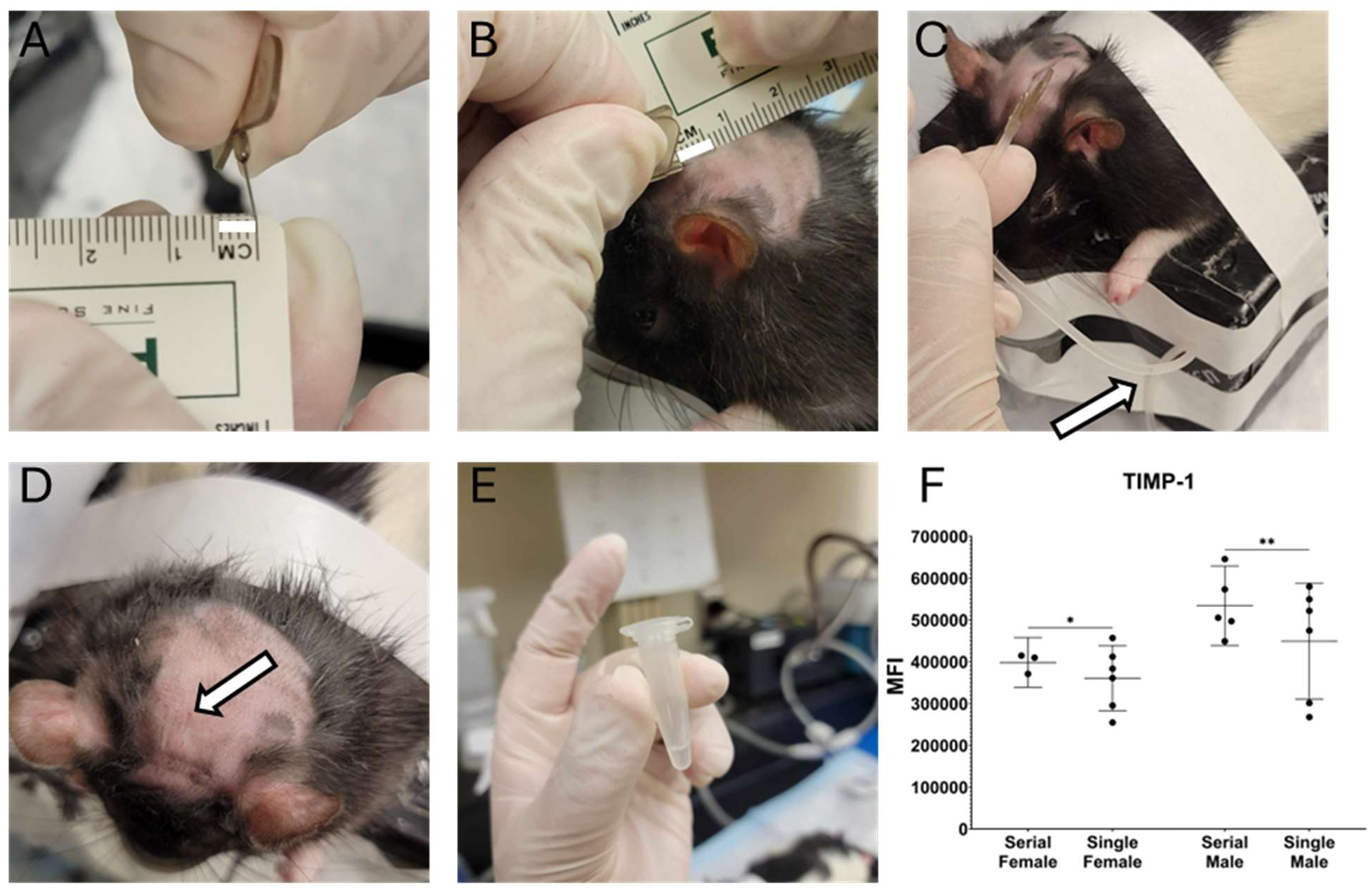
CSF withdrawal procedure. First, a 27G winged needle tip is bent approximately 90° with a 4 mm tip (emphasized by the white rectangle) (A). The needle is inserted into the foramen magnum, an extremely soft spot approximately 6 mm (emphasized by the white rectangle) past the external occipital proturberance of the skull (B). Clear CSF (white arrow) should flow into the tubing of the winged needle (C). After removal of the needle, there should not be any visible signs of damage on the skin, aside from the insertion point (white arrow) (D). On average, 90-100 μL of clear CSF can be collected (E). Our statistics show that collection frequency does not affect the expression of any analyte on the panel, except TIMP1, in males and females (F).

### 3.2. Serial CSF collection did not alter protein detection

Serially collected 7dpi CSF samples were compared to CSF samples that were collected only at 7 dpi. Analysis by two-way ANOVA demonstrated that collection frequency did not alter protein levels in either males (F=2.492; *p*=0.1135) or females (F=1.022; *p=*0.3140) across 16 of the 17 total protein analytes. Interestingly, TIMP-1 was significantly increased (*p*=0.0143 in females and *p*=0.0055 in males) in serially collected CSF samples using a post-hoc Holm-Šídák’s multiple comparisons test (Figure 2F). TIMP-1 is an inhibitor of matrix metalloproteinases, and has been shown to play a role in alleviating inflammatory pain associated with wound healing^33^. Given the role of TIMP-1 role in inflammatory pain and wound healing, this result is consistent with specific modulation of this pathway. Together, these data suggest that while detection of protein changes over time is possible, repeated CSF collection did not result in a generalized alteration of protein analyte detection.

### 3.3. 16 of 17 CSF protein analytes were unaltered by a single mTBI when data were analyzed without consideration of sex

To interrogate the sex differences in post-concussion CSF protein levels, we took a progressive approach to analyzing the data. First, we compared CSF protein levels between concussion and sham to assess how a single concussion alters the proteome of the CSF without consideration for sex. Several publications presenting clinical concussion data ignore sex, often grouping patients into two groups: concussion vs sham^9,16,34–36^. Using this approach, we compared concussion and sham CSF samples normalized to pre-injury levels using two-way mixed effects analysis and post-hoc Holm-Šídák’s multiple comparisons test (Table 1). A single concussion had no significant effect on any of the 17 analytes, except for interferon-γ (IFNγ). For IFNγ (Figure 3), both time (F=14.46; *p*<0.0001) and injury (F=5.408; *p=*0.0270) had significant effects, however the interaction (F=1.023; *p*=0.3891) of these two factors had no significant effect.

**Table 1:** Statistical testing comparing injury to sham, ignoring sex. Two-way mixed effects analysis with post-hoc Holm-Sidak multiple comparisons test. Ignoring sex, analyte mean fluorescent intensities (MFI) at each time point were normalized to pre-injury measurements and compared between concussion and sham. \**p*<0.05; \*\*\*\**p*<0.0001

| Sex not considered | Two-way Mixed Effects Analysis |  |  | Post-hoc Holm-Sidak Multiple Comparisons (Concussion vs Sham) |  |  |  |
| --- | --- | --- | --- | --- | --- | --- | --- |
|  | Time | Injury | Time x Injury | 1dpi | 3 dpi | 7 dpi | 14 dpi |
| Protein |  |  |  |  |  |  |  |
| CXCL3 | ns | ns | ns | ns | ns | ns | ns |
| CXCL2 | ns | ns | ns | ns | ns | ns | ns |
| GM-CSF | ns | ns | ns | ns | ns | ns | ns |
| ICAM-1 | ns | ns | ns | ns | ns | ns | ns |
| IFNg | **** | * | ns | ns | ns | ns | ns |
| IL-1a | ns | ns | ns | ns | ns | ns | ns |
| IL-1b | ns | ns | ns | ns | ns | ns | ns |
| IL-2 | ns | ns | ns | ns | ns | ns | ns |
| IL-4 | ns | ns | ns | ns | ns | ns | ns |
| IL-6 | ns | ns | ns | ns | ns | ns | ns |
| IL-10 | ns | ns | ns | ns | ns | ns | ns |
| IL-13 | ns | ns | ns | ns | ns | ns | ns |
| IL-18 | ns | ns | ns | ns | ns | ns | ns |
| L-selectin | ns | ns | ns | ns | ns | ns | ns |
| TIMP-1 | ns | ns | ns | ns | ns | ns | ns |
| TNFa | ns | ns | ns | ns | ns | ns | ns |
| VEGF | ns | ns | ns | ns | ns | ns | ns |

**Figure 3:**
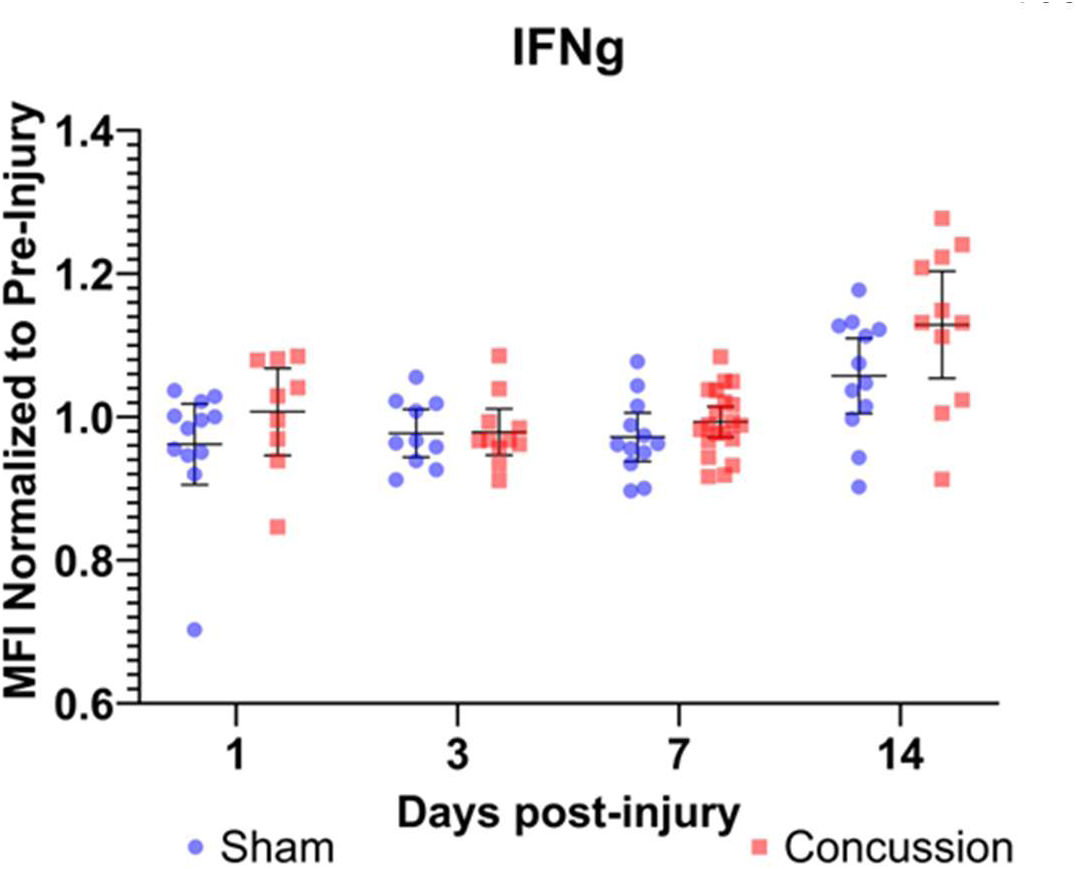
IFNg increases slightly over time after concussion. Graph shows the mean fluorescent intensity (MFI) at each timepoint normalized to pre­-injury. Mean with 95% Cl is plotted with individual points representing Individual rats.

### 3.4. Males exhibit a greater inflammatory response to a single concussion than females

We next separated samples by sex, comparing concussion and sham animals within each. Using the same statistical approach as before, females showed significant responses to injury in multiple pro-resolving/anti-inflammatory proteins, whereas males showed significant responses to injury in several pro-inflammatory proteins. For females, two-way Mixed Effects analysis (Table 2) identified a significant interaction effect for time and injury on IL-13 (F=3.066, *p=*0.0388, Figure 4A), however, neither time (F=0.06031, *p*=0.9103) nor injury (F=0.001776, *p*=0.9666) alone had a significant effect on IL-13 levels. Similarly, VEGF (Figure 4C) showed comparable results, where the interaction of time and injury (F=5.436, *p*=0.0049) had a significant effect on VEGF levels, but neither time (F=2.164, *p*=0.1352) nor injury (F=0.006961, *p*=0.9348) alone had significant effects on VEGF levels. Third, time (F=8.068, *p*=0.0065) and injury (F=7.458, *p*=0.0095) both affected the levels of soluble L-selectin, an adhesion protein found in leukocytes, (Figure 4B) in injured female rats but not the interaction of time and injury (F=1.267, *p*=0.2995).

**Table 2:** Statistical testing comparing injury to sham in females. Two-way mixed effects analysis with post-hoc Holm-Sidak multiple comparisons test comparing concussion vs sham in female CSF samples only. Analyte mean fluorescent intensities (MFI) at each time point were normalized to pre-injury measurements and compared between concussion and sham. \**p*<0.05; \*\**p*<0.01

| Females Only | Two-way Mixed Effects Analysis |  |  | Post-hoc Holm-Sidak Multiple Comparisons<br>Males (Concussion vs Sham) |  |  |  |
| --- | --- | --- | --- | --- | --- | --- | --- |
| Protein | Time | Injury | Time x Injury | 1dpi | 3 dpi | 7 dpi | 14 dpi |
| CXCL3 | ns | ns | ns | ns | ns | ns | ns |
| CXCL2 | ns | ns | ns | ns | ns | ns | ns |
| GM-CSF | * | ns | ns | ns | ns | ns | ns |
| ICAM-1 | ns | ns | ns | ns | ns | ns | ns |
| IFNg | ** | ns | ns | ns | ns | ns | ns |
| IL-1a | ns | ns | ns | ns | ns | ns | ns |
| IL-1b | * | ns | ns | ns | ns | ns | ns |
| IL-2 | ns | ns | ns | ns | ns | ns | ns |
| IL-4 | ns | ns | ns | ns | ns | ns | ns |
| IL-6 | * | ns | ns | ns | ns | ns | ns |
| IL-10 | ns | ns | ns | ns | ns | ns | ns |
| IL-13 | ns | ns | * | ns | ns | ns | ns |
| IL-18 | ns | ns | ns | ns | ns | ns | ns |
| L-selectin | ** | ** | ns | ns | ns | ns | ns |
| TIMP-1 | ns | ns | ns | ns | ns | ns | ns |
| TNFa | ns | ns | ns | ns | ns | ns | ns |
| VEGF | ns | ns | ** | ns | ns | ns | ns |

**Figure 4:**
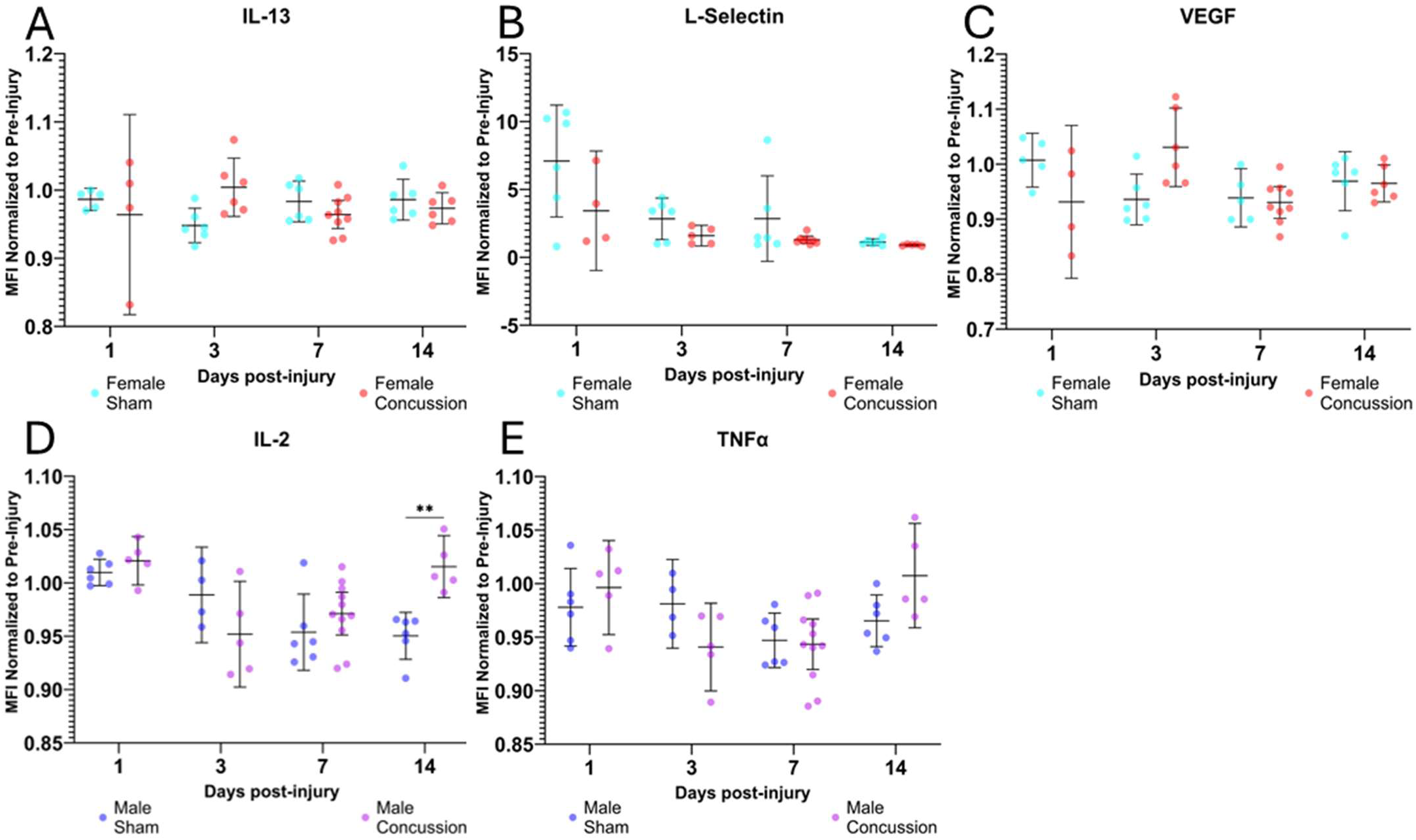
Injury is a significant factor affecting expression in pro-resolving proteins forfemales and pro-inflammatory proteins for males. The MFI for each sample at each timepoint normalized to pre-injury is reported for IL-13 (A), L-Selectin (B), and VEGF (C) in females, as well as IL-2 (D) and TNFα (E) in males. These proteins were selected based on results from statistical testing reported in Table 2 and 3. Mean with 95% Cl is plotted with individual points representing individual rats. \*\**p*<0.01.

For males, two pro-inflammatory proteins exhibited clear modulation (Table 3). First, a two-way Mixed Effects analysis reported time (F=8.598; p=0.0005) and the interaction of time and injury (F=5.732; p=0.0023) to be significant factors affecting interleukin-2 (IL-2, Figure 4D) levels with a significant increase in IL-2 levels after concussion at 14 dpi (*p*=0.0051) in male rats. Injury alone was not shown to be a significant factor affecting IL-2 levels over time (F=2.955; *p*=0.0933).

**Table 3:** Statistical testing comparing injury to sham in males. Two-way mixed erects analysis with post-hoc Holm-Sidak multiple comparisons test comparing concussion vs sham in male CSF samples only. Analyte mean fluorescent intensities (MFI) at each time point were normalized to pre-injury measurements and compared between concussion and sham. \**p*<0.05; \*\**p*<0.01; \*\*\**p*<0.001

| Males Only | Two-way Mixed Effects Analysis |  |  | Post-hoc Holm-Sidak Multiple Comparisons<br>Males (Concussion vs Sham) |  |  |  |
| --- | --- | --- | --- | --- | --- | --- | --- |
| Protein | Time | Injury | Time x Injury | 1dpi | 3 dpi | 7 dpi | 14 dpi |
| CXCL3 | ns | ns | ns | ns | ns | ns | ns |
| CXCL2 | ns | ns | ns | ns | ns | ns | ns |
| GM-CSF | ns | ns | ns | ns | ns | ns | ns |
| ICAM-1 | ns | ns | ns | ns | ns | ns | ns |
| IFNg | * | ns | ns | ns | ns | ns | ns |
| IL-1a | ns | ns | ns | ns | ns | ns | ns |
| IL-1b | ns | ns | ns | ns | ns | ns | ns |
| IL-2 | *** | ns | ** | ns | ns | ns | ** (S < C) |
| IL-4 | ns | ns | ns | ns | ns | ns | ns |
| IL-6 | ns | ns | ns | ns | ns | ns | ns |
| IL-10 | ns | ns | ns | ns | ns | ns | ns |
| IL-13 | ns | ns | ns | ns | ns | ns | ns |
| IL-18 | ** | ns | ns | ns | ns | ns | ns |
| L-selectin | ** | ns | ns | ns | ns | ns | ns |
| TIMP-1 | ns | ns | ns | ns | ns | ns | ns |
| TNFa | ** | ns | * | ns | ns | ns | ns |
| VEGF | ** | ns | ns | ns | ns | ns | ns |

Second, a two-way Mixed Effects analysis reported time (F=5.549; p=0.0067) and the interaction of time and injury (F=3.185; p=0.0412) to be significant factors affecting tumor necrosis factor-α (TNFα, Figure 4E) levels. Injury alone was not shown to be a significant factor affecting TNFα levels (F=0.07991; *p*=0.7813), and there were no significant pairwise comparisons between sham and concussion at any specific timepoints.

Interestingly, both males and females had several proteins give time alone as a significant factor affecting protein levels. In females (Table 2), time was significant for GM-CSF (F=3.939, *p*=0.0298), IFNγ (F=7.900, *p*=0.0065), IL-1β (F=5.653, *p*=0.0109), and IL-6 (F=3.765, *p*=0.0462). L-Selectin (F=6.145, *p*=0.0056). In males (Table 3), time was significant for IFNγ (F=7.555, *p*=0.0111), IL-18 (F=9.450, *p*=0.0022), L-selectin (F=7.434, p=0.0094), and VEGF (F=3.733, *p*=0.0413).

### 3.5. Sex is a significant factor in CSF protein levels after a single concussion, with males showing significantly greater levels of pro-inflammatory proteins up to 7 dpi

Third, we compared the CSF protein levels of males and females in response to a single concussion (Table 4). Sex was found to be a significant factor affecting the levels of 9 proteins: CXCL3 (F=11.05, *p*=0.0019, Figure 5A), GM-CSF (F=7.604; *p*=0.0085, Figure 5B), ICAM-1 (F=6.451, *p=*0.0211, Figure 5C), IL-1β (F=8.072, *p*=0.0108, Figure 5D), IL-10 (F=6.006; *p*=0.0186, Figure 5G), IL-18 (F=5.137; *p*=0.0285, Figure 5H), L-selectin (F=5.585, *p*=0.0303, Figure 5I), TNFα (F=4.083, *p*=0.0496, Figure 5J), and VEGF (F=4.704, *p*=0.0437, Figure 5K). Additionally, time was also found to be a significant factor affecting the levels of IFNγ (F=10.45, *p*=0.0012), IL-18 (F=5.164, *p*=0.0225), and L-selectin (F=6.101, *p*=0.0192). Further, the interaction of sex and time was shown to significantly affect the levels of an additional three proteins: IL-1b (F=3.531; *p*=0.0300, Figure 5D), IL-2 (F=3.683; *p*=0.0191, Figure 5E), and VEGF (F=7.095; *p*=0.0014, Figure 5K). Post-hoc multiple comparisons testing comparing CSF protein levels after a single concussion in males vs females at specific timepoints showed no significant differences at 1 dpi and 14 dpi. However, at 3 dpi females had significantly higher TNFα levels than males (*p*=0.0275, Figure 5J). At 7 dpi, males showed significantly greater levels of three proteins compared to females: IL-6 (*p*=0.0200, Figure 5F), IL-18 (*p*=0.0410, Figure 5H), and VEGF (*p*=0.0113, Figure 5K).

**Table 4:**
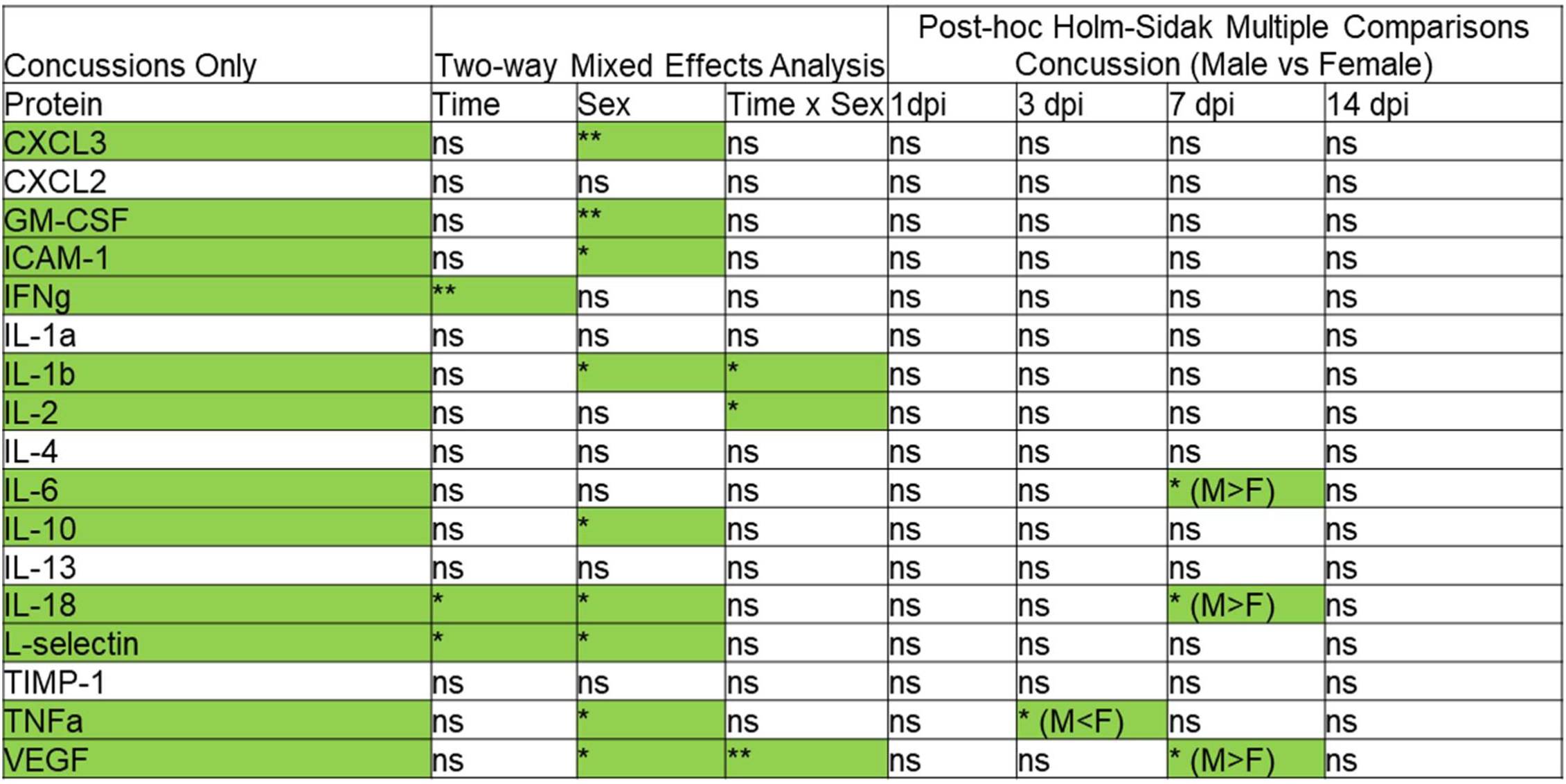
Statistical testing comparing injured males to injured females. Two-way mixed effects analysis with post-hoc Holm-Sidak multiple comparisons test comparing male vs female in concussion CSF samples only. Analyte mean fluorescent intensities (MFI) at each time point were normalized to pre-injury measurements and compared between male and female. \**p*<0.05; \*\**p*<0.01

| Concussions Only | Two-way Mixed Effects Analysis |  |  | Post-hoc Holm-Sidak Multiple Comparisons<br>Concussion (Male vs Female) |  |  |  |
| --- | --- | --- | --- | --- | --- | --- | --- |
|  | Time | Sex | Time x Sex | 1dpi | 3 dpi | 7 dpi | 14 dpi |
| Protein |  |  |  |  |  |  |  |
| CXCL3 | ns | ** | ns | ns | ns | ns | ns |
| CXCL2 | ns | ns | ns | ns | ns | ns | ns |
| GM-CSF | ns | ** | ns | ns | ns | ns | ns |
| ICAM-1 | ns | * | ns | ns | ns | ns | ns |
| IFNg | ** | ns | ns | ns | ns | ns | ns |
| IL-1a | ns | ns | ns | ns | ns | ns | ns |
| IL-1b | ns | * | * | ns | ns | ns | ns |
| IL-2 | ns | ns | * | ns | ns | ns | ns |
| IL-4 | ns | ns | ns | ns | ns | ns | ns |
| IL-6 | ns | ns | ns | ns | ns | * (M>F) | ns |
| IL-10 | ns | * | ns | ns | ns | ns | ns |
| IL-13 | ns | ns | ns | ns | ns | ns | ns |
| IL-18 | * | * | ns | ns | ns | * (M>F) | ns |
| L-selectin | * | * | ns | ns | ns | ns | ns |
| TIMP-1 | ns | ns | ns | ns | ns | ns | ns |
| TNFa | ns | * | ns | ns | * (M<F) | ns | ns |
| VEGF | ns | * | ** | ns | ns | * (M>F) | ns |

**Figure 5:**
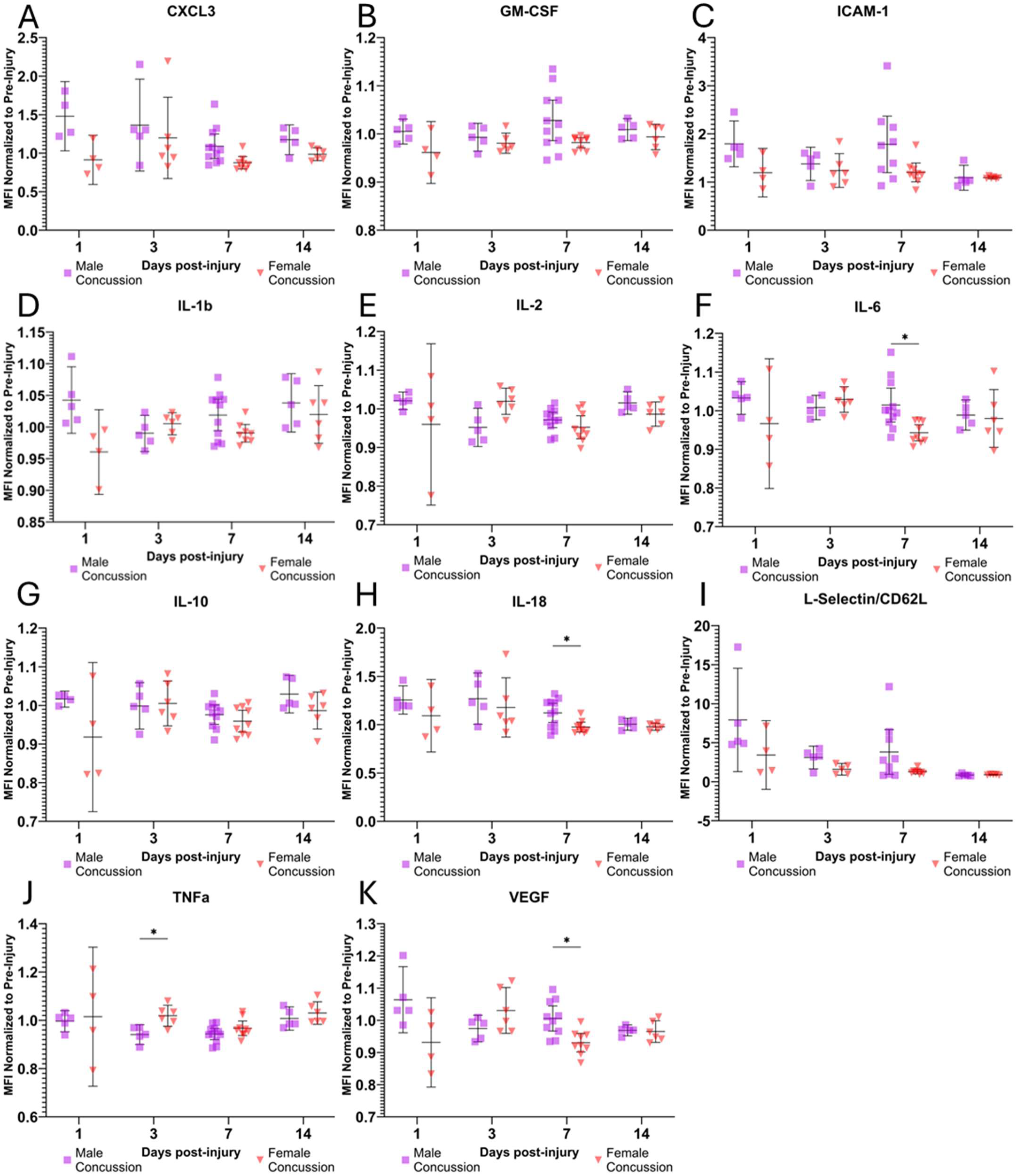
Males show a delayed inflammatory response to concussion compared to females. The MFI for each sample at each timepoint normalized to pre-injury is reported forCXCL3 (A), GM-CSF (B), ICAM-1 (C), IL-1 b (D), IL-2 (E), IL-6 (F), IL-10 (G), IL-18 (H), L-selectin/CD62L (I), TNFα (J), and VEGF (K). These proteins were selected based on results from statistical testing reported in Table 4 showing sex as a significant factor affecting protein expression. Mean with 95% Cl is plotted with individual points representing individual rats. \**p*<0.05.

### 3.6. Male and female sham animals exhibit baseline differences in CSF protein detection

The last component of our CSF analysis was comparing all four groups: Male concussion, male sham, female concussion, and female sham in a combined model. Post-hoc multiple comparisons testing (Table 5) showed significant differences when comparing sham male and sham female to groups of the opposite sex. At 1 and 14 dpi, there were no significant differences in the levels of any of the 17 proteins when comparing either sham male or sham female with the opposite sex. However, at 3 dpi, sham male was shown to have significantly greater levels of CXCL3 (*p*=0.0030, Figure 6A) and VEGF (*p*=0.0145, Figure 6G) than sham female. At 7dpi, sham male was shown to have significantly greater levels of CXCL3 (*p*=0.0339, Figure 6A) and CXCL2 (*p*=0.0404, Figure 6B), as well as significantly lower levels of IFNγ (Figure 6C) compared to concussed female (*p*=0.0181) and sham female (*p*=0.0489). Additionally, using this statistical approach, which includes the most comparisons of all statistical tests, and thus the least statistical power in this study, 4 of the 5 significant pairwise comparisons reported in the previous tests were confirmed (Table 5) at this lower power. At 3 dpi, IFNγ was significantly greater in concussed females compared to concussed males (*p*=0.0409, Figure 6C). At 7 dpi, concussed males had significantly greater levels of IL-6 (*p*=0.0298, Figure 6E) and VEGF (*p=*0.0169, Figure 6G) compared to concussed females. Additionally, at 7 dpi, we previously reported the concussed males had significantly greater levels of IL-18 (Figure 5H), but in this lower powered analysis, this comparison is no longer significant (*p*=0.0608). At 14 dpi, concussed males had significantly greater levels of IL-2 (*p*=0.0076, Figure 6D) compared to uninjured males.

**Table 5:** Statistical testing comparing all four experimental groups. Two-way mixed effects analysis with post-hoc Holm-Sidak multiple comparisons test comparing all 4 experimental groups. Analyte mean fluorescent intensities (MFI) at each time point were normalized to pre-injury measurements and compared between groups. MS = Male Sham, MC = Male Concussion, FS = Female Sham, FC = Female Concussion \**p*<0.05; \*\**p*<0.01.

| All 4 groups | Post-hoc Holm-Sidak Multiple Comparisons Concussion (Male vs Female) |  |  |  |
| --- | --- | --- | --- | --- |
| Protein | 1dpi | 3 dpi | 7 dpi | 14 dpi |
| CXCL3 | ns | ** (MS>FS) | * (MS>FC) | ns |
| CXCL2 | ns | ns | * (MS>FC) | ns |
| GM-CSF | ns | ns | ns | ns |
| ICAM-1 | ns | ns | ns | ns |
| IFNγ | ns | ns | * (MS<FS); * (MS<FC) | ns |
| IL-1a | ns | ns | ns | ns |
| IL-1b | ns | ns | ns | ns |
| IL-2 | ns | ns | ns | ** (MS<MC) |
| IL-4 | ns | ns | ns | ns |
| IL-6 | ns | ns | * (MC>FC) | ns |
| IL-10 | ns | ns | ns | ns |
| IL-13 | ns | ns | ns | ns |
| IL-18 | ns | ns | ns | ns |
| L-selectin | ns | ns | ns | ns |
| TIMP-1 | ns | ns | ns | ns |
| TNFα | ns | * (MC<FC) | ns | ns |
| VEGF | ns | * (MS>FS) | * (MC>FC) | ns |

**Figure 6:**
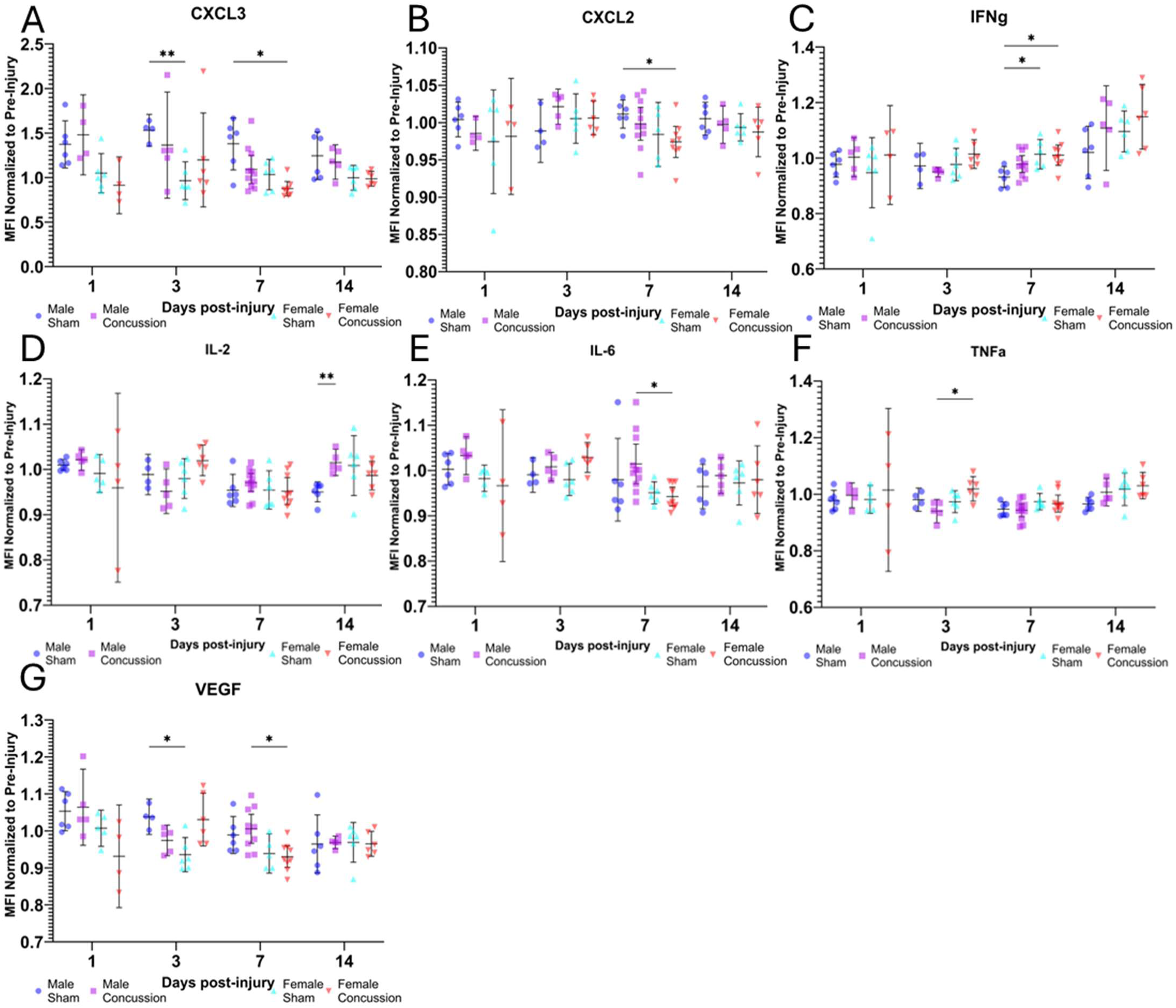
Males and females have varying levels of analytes in sham groups which can confound analysis. The MFI for each sample at each timepoint normalized to pre-injury is reported for CXCL3 (A), CXCL2 (B), IFNg (C), IL-2 (D), IL-6 (E), TNFα (F), and VEGF (G). These proteins were selected based on results from statistical testing reported in Table 5 showing significant pairwise comparisons. Mean with 95% Cl is plotted with individual points representing individual rats. *p<0.05, **p<0.01

### 3.7. Males exhibit changes in blood-brain-barrier (BBB) permeability and microglia number after a concussion, but not females

In addition to analyzing changes in CSF protein levels, we performed histological characterization of the brain tissue. We quantified the numbers of astrocytes (GFAP+, Figure S1) and microglia (Iba1+, Figure 7) using stereological counting and performed a two-way ANOVA and post-hoc Holm-Šídák’s multiple comparisons test on the estimated number of astrocytes. Our two-way ANOVA showed that neither injury (*p*=0.6664, F=0.4115), sex (*p*=0.8045, F=0.06237), nor the interaction of the two (*p*=0.9444, F=0.05730) were significant factors affecting the number of GFAP+ astrocytes. Conversely, our two-way ANOVA showed that injury (*p*=0.0346, F=3.772) significantly affected the number of Iba1+ microglia. Post-hoc multiple comparisons testing reveals a significant increase (*p*=0.0151) in Iba1+ microglia from 7 dpi to 14 dpi in males (Figure 7G). Neither sex (*p*=0.3142, F=1.048) nor the interaction of sex and injury (*p=*0.3022, F=1.246) significantly affected the number of microglia. While not significant, there is a slight decrease in the number of microglia at 7 dpi compared to sham in males. There were no significant changes in the number of microglia in females, but at 14 dpi, the number of microglia seems to be diverging between males and females (Figure 7G).

**Figure 7:**
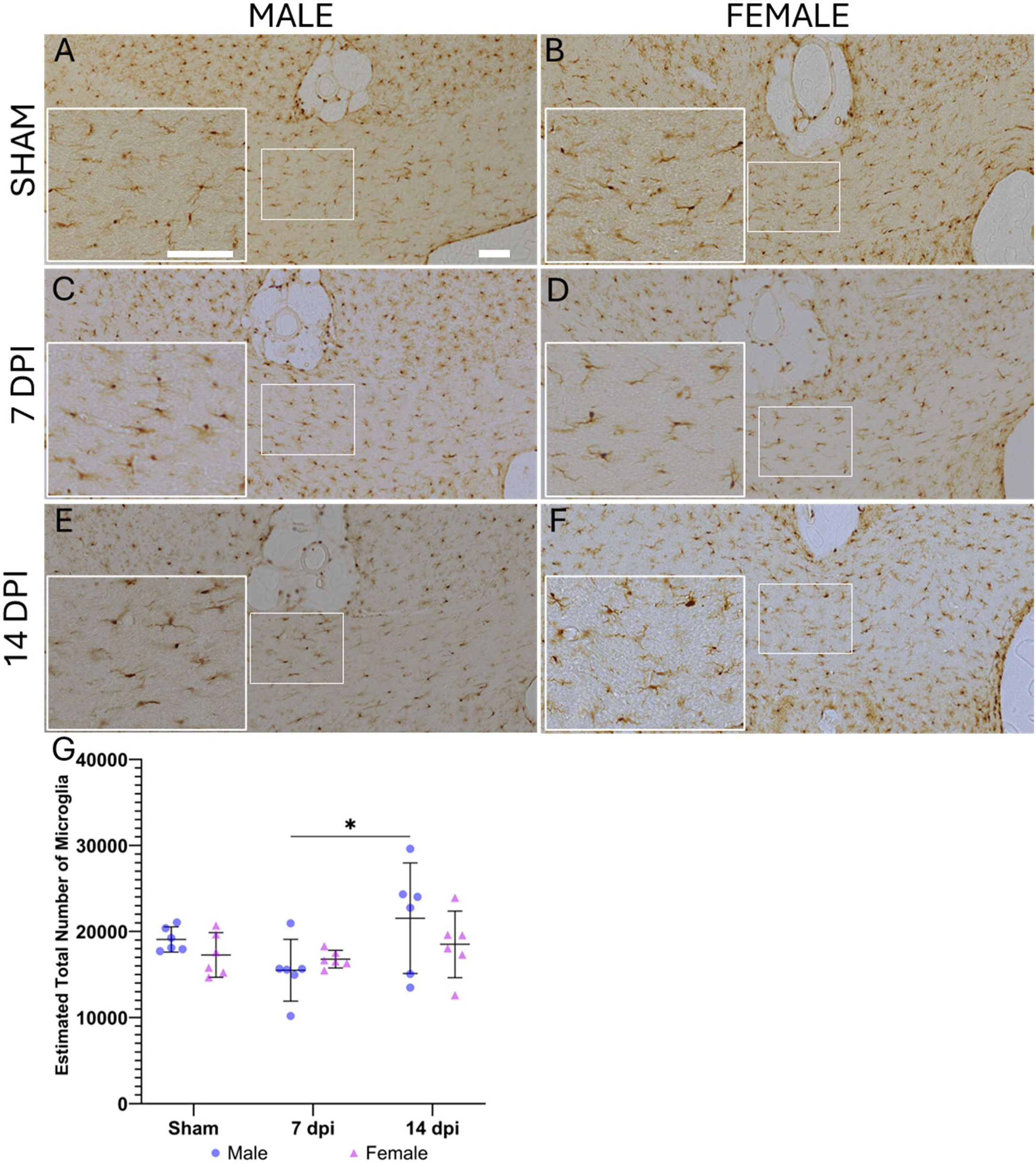
Males experience a significant increase in the estimated total number of microglia, but not females, 14 days after a concussion. Stereological quantification of Iba1 + cells was performed on serial sections of brain tissue. Representative images are shown (A-F). Quantification was performed only in the corpus callosum and is reported as the estimated total number of microglia (G). Data is presented as mean with 95% Cl. Statistical analysis was performed using a two-way ANOVA and post-hoc Holm-Sidak multiple comparisons test. *p*<0.05. Scale bar = 100 μm.

The trends in this inflammatory cell data correspond well with trends seen in scoring of Prussian Blue staining for microbleeds (Figure 8). Microbleeds were semi-quantitatively analyzed as previously mentioned. While two-way ANOVA revealed that sex (*p*=0.0552, F=3.981), injury (*p*=0.0749, F=2.829), or the interaction of the sex and injury (*p*=0.0511, F=3.289) did not significantly affect the presence of microbleeds, post-hoc Holm-Šídák’s multiple comparisons testing shows there are significant differences between groups (Figure 8G). In males, there is a significantly greater presence of microbleeds at 14 dpi compared to sham (*p=*0.0376) and 7 dpi (*p=*0.0073). Further, at 14 dpi there is a significantly greater presence of microbleeds in males compared to females (*p*=0.0029). There are no significant differences in the presence of microbleeds in females. Further, co-staining of Iba1 and Prussian Blue reveal microglial phagocytosis of microbleeds (Figure 9).

**Figure 8:**
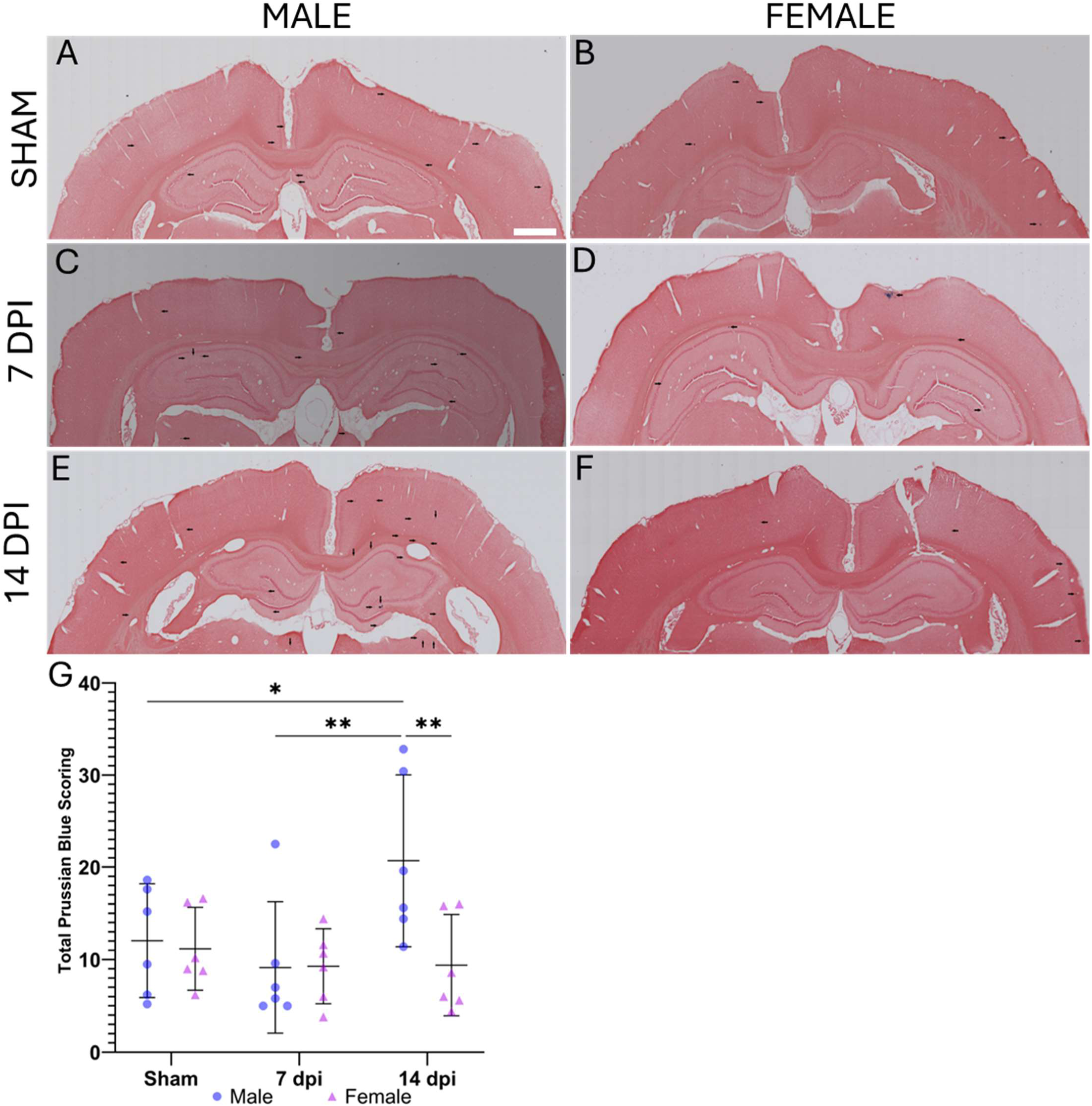
Males, but not females, experience a significant increase in the presence of microbleeds 14 days after a concussion. A semi-quantitative scoring system for microbleed presence in the cortex, corpus callosum, and hippocampus was used. Representative images are shown (A-F). Black arrows, added by someone blinded to the experimental groups, signify microbleeds in the images. The total scores of 5 brain sections were summed up for each animal and used for statistical testing (G). Data is presented as mean with 95% Cl. Statistical analysis was performed using a two­way ANOVA and post-hoc Holm-Sidak multiple comparisons test. \**p*<0.05. \*\**p*<0.01 Scale bar = 1000 μm.

**Figure 9:**
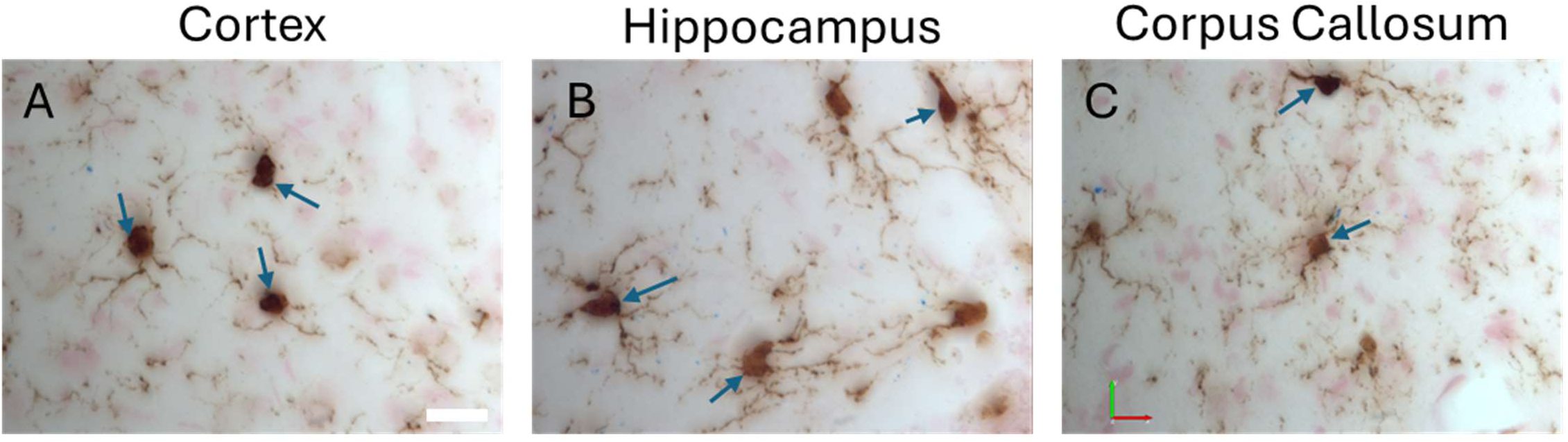
Microglia phagocytose microbleeds. Representative images show colocalized positive staining for Iba1 and microbleeds confirming microgliaare responding to BBB permeability in the cortex (A), hippocampus(B), and corpus callosum (C). Scale bar = 20 μm.

## 4. Discussion

While mTBI has been widely studied, there are still many aspects of the pathophysiology that remain poorly understood. First, it is unclear if repeated collection of CSF from the same animals will alter the proteome being investigated and confound the results. Second, it has been quite challenging to fully understand the temporal dynamics of the neuroinflammatory response to mTBI as currently established methods (i.e. stand-alone CSF collection, brain tissue lysates, histology) require large numbers of animals for studies. Third, the majority of mTBI literature fails to investigate sex as a biological variable. This study addressed these limitations.

### 4.1. Serial collection of CSF does not significantly alter CSF inflammation

We first described a method for minimally invasive serial collection of CSF. CSF proteomics are not widely studied in pre-clinical models due to the difficult nature of CSF collection. Previously described methods of CSF collection do not allow for repeated collections, require specialized tools and equipment, and can cause further brain injury or death^26,27,37^. The serial collection method used herein did not cause the rats to experience unusually significant weight changes, exhibit evidence of pain/distress (shallow or labored breathing, poor grooming, hunching, or porphyria), show signs of infection or swelling, or die.

Further, since serial collection of CSF requires multiple punctures with a needle over time, we compared a 7 dpi fresh CSF sample from rats who underwent serial collection at pre-injury, 1 dpi, 3 dpi, 7 dpi, and 14 dpi to a CSF sample from rats who only underwent a single CSF collection at 7 dpi (Figure 1). A two-way ANOVA revealed that collection frequency was not a significant factor affecting the proteome results, at least for the 17 analytes we assessed. Interestingly, however, a post-hoc Holm-Šídák’s multiple comparisons test revealed that none of the analytes in our panel showed significant differences between serial collection and single collection except TIMP-1 in both males and females (Figure 2). TIMP-1 is an inhibitor of metalloproteinases that plays a role in alleviating inflammatory pain associated with wound healing^33^. Given this role for TIMP-1, the elevation in TIMP-1 resulting from repeated collection of CSF makes sense as repeated punctures would cause increased inflammation and prolonged wound healing. This one elevated protein provides confidence in our statistical testing as it shows that the statistical approach has enough power to detect a significant result and supports the assumption that our negative statistical results using similar methods are unlikely to be false negatives.

The establishment of an easily adapted serial CSF collection method is also critical to closing the translational gap as it enables more longitudinal studies to be performed with a clinically relevant output. Clinically relevant outputs used in longitudinal studies can help address the problem of translating time between rodents and humans. Such a translation of time scale is inherently difficult, as there is no universal conversion factor, presenting a major obstacle in relating preclinical studies to clinical studies.

### 4.2. In sex-blended analysis, a single concussion only affects the levels of IFNγ in the CSF

After showing our method of serial CSF collection is not globally increasing inflammatory protein levels in the CSF, we began to interrogate the effects of a single concussion on inflammation and the potential sex-based differences in analytes from CSF. To do this, we took a stepwise approach to analyzing the levels of proteins in the CSF. First, we asked how CSF protein levels are affected by a single concussion, without considering sex. Many clinical mTBI studies do not consider sex, combining male and female samples, only comparing injured and uninjured groups^9,16,34–36^. This is often due to insufficient numbers of female patient samples. For our statistical test, we combined male and female concussion groups and compared them to combined male and female sham groups, effectively giving us two groups: concussion vs sham. The results from this test were rather interesting, as there were not any significant differences in protein levels over time (Table 1), with the exception of IFNγ. IFNγ (Figure 3) was shown to be significantly affected by injury.

IFNγ has been widely studied as a biomarker for mTBI, however there is little agreement across studies due to a lack of uniformity in study designs. In sex blended studies, IFNγ has been shown to be significantly increased acutely after mTBI in blood samples from children^14^ and adults^36^. Further, in a sex blended longitudinal study collecting blood samples from adult mTBI patients at admission to 12 months post injury by Chaban and colleagues, IFNγ was shown to be significantly affected by injury in a two-way mixed effects analysis^38^. While this result from blood testing directly reflects our finding in this study using CSF, it is important to note that Chaban *et al*. did not report time as a significant factor influencing IFNγ levels. Interestingly, adult patients showed no significant differences in IFNγ levels compared to controls in CSF samples taken approximately 17 months after concussion^9^. It should be noted that the athletes in this sample had a median of 5 concussions prior to CSF collection.

While IFNγ was the only protein in our panel that showed significance in our sex blended statistical testing, TNFα, IL-2, IL-6, IL-10, and VEGF are also interesting, as these five proteins have shown significant changes in sex blended human studies. Chaban *et al.* reported that TNFα expression in adult mTBI patient blood samples was significantly affected by both time and injury^38^. The 207 patients in this study sustained a single mild TBI, but the injury was severe enough to result in hospitalization. Vedantam *et al.* showed that IL-2 and IL-6 are both significantly increased compared to controls at 1 dpi and IL-6 stayed significantly increased up to 6 months post-injury in blood samples from adult mTBI patients, and these changes were associated with more severe symptoms, depression, and/or PTSD. Again, these patients were recruited from injuries serious enough to warrant a trip to the emergency room^36^. Ryan *et al*. reported that TNFα and IL-10 were significantly decreased, while IL-6 was significantly increased in blood sampled from pediatric mTBI patients within 2 weeks of injury^14^. Gard *et al*. reported that IL-2, TNFα, and VEGF were all significantly increased in the CSF of adult mTBI patients 17 months post-injury^9^. While our results do not fully reflect what has been published in sex blended clinical studies, it is important to note that while these studies are sex blended, they are still predominantly male patient populations with almost a 2:1 male:female ratio^9,14,36,38^. To fully understand these published results, as well as our results in this study, it is vital to separate the sexes and compare the injury response in each sex.

### 4.3. A single concussion significantly affects pro-resolving proteins in females and pro-inflammatory proteins in males

The second step we took in our stepwise approach to analyzing this dataset was separating the sexes and comparing the injury response in each sex. This approach gave us two sets of two groups: Concussion vs sham in females (Table 2) and concussion vs sham in males (Table 3). The results from this approach were quite intriguing. In females, injury was a significant factor affecting the levels of soluble L-selectin (Figure 4B) and the interaction of time and injury was a significant factor affecting the levels of IL-13 (Figure 4A) and VEGF (Figure 4C). In males, the interaction of time and injury significantly affected the levels of IL-2 (Figure 4D) and TNFα (Figure 4E). IL-13 and VEGF are typically secreted by pro-resolving, anti-inflammatory immune cells, while IL-2 and TNFα are pro-inflammatory proteins^39–41^. The trends shown in the data paint an interesting picture. Females experience an acute upregulation of pro-resolving proteins, which have also been shown to be neuroprotective^41,42^, while males initially experience a decrease in pro-inflammatory proteins that eventually increases at later timepoints, with IL-2 levels being significantly greater in concussed males compared to sham males at 14 dpi, which reflects previously published male-biased sex blended studies^9,36^. One possible explanation of this paradigm is the differential presence of estrogen and progesterone, which have been shown to be neuroprotective after TBI, reducing mortality and improving neurological outcomes, by reducing neuronal apoptosis, inhibiting microglial and astrocytic activation, and decreasing brain edema^17,20–22^. To further dissect the sex differences in the inflammatory response after concussion, it is imperative to compare CSF protein levels after concussion across both sexes directly.

### 4.4. Males experience a biphasic and more prolonged inflammatory response compared to females

The third phase of our stepwise approach to analyzing CSF protein concentration after concussion was comparing the protein levels in both concussion groups, giving us two groups for comparison: male vs female. In this comparison, many sex-based differences were revealed. Sex was a factor significantly affecting the levels of 9 of 17 proteins and the interaction of sex and time was a factor significantly affecting the levels of 3 of these proteins (Table 4). There is a clear trend present in the time course representations of each of these proteins (Figure 5). While there are only four significant pairwise comparisons in post-hoc multiple comparisons testing, in general, males experience a more rapid inflammatory response, with all proteins being greater in males than females, except TNFα, at 1 dpi. At 3 dpi, females have significantly greater TNFα than males, as well as greater levels of IL-1β, IL-2, IL-6, and VEGF, suggesting that females undergo a delayed inflammatory response. At 7 dpi, males experience a second peak in inflammation with significantly greater levels of IL-6, IL-18, and VEGF, as well as greater levels of CXCL3, GM-CSF, ICAM-1, IL-1b, IL-2, and L-selectin, than females. This persists at 14 dpi, with males exhibiting greater levels of CXCL3, GM-CSF, IL-1b, IL-2, and IL-10, though none of these are significant, than females.

Our results are supported by a previous study evaluating serum and brain cytokine concentrations in mice 4 hours after the last injury in a repetitive blast-related mTBI model^28^. Baskin *et al.* reported that female mice experience greater anti-inflammatory signaling acutely after mTBI, with significantly higher concentrations of IL-10 and G-CSF and significantly lower concentrations of IL-9, IL-12, and IP-10 in brain tissue homogenates, compared to male mice^28^. While few studies, especially longitudinal ones, have compared protein levels in biofluids sampled from mTBI patients across sexes, it is critical to analyze these sex differences more holistically, as biomarker bioactivity may vary between sexes. For example, Di Battista *et al.* reported no significant differences in plasma concentrations of IFNγ or TNFα between male and female mTBI patients^43^. However, in males, they found that higher IFNγ concentrations in blood were significantly correlated with more severe concussion symptoms, whereas in females, higher concentrations of IFNγ and TNFα were significantly correlated with less severe symptoms^43^. This data suggests there are underlying differences in the mechanisms regulating how males and females respond to inflammation. Studies comparing male and female injury responses to mTBI are valuable for bridging the sex-based knowledge gap, and more studies investigating these sex-based differences are clearly needed. With that said, directly comparing the injuries between sexes is not the complete picture. It is imperative to understand uninjured baseline differences between sexes as well, especially in longitudinal biomarker studies such as the present one.

### 4.5. Uninjured males and females have varying cytokine levels in the CSF

The last phase in our stepwise approach to analyzing this dataset incorporates the sham groups into the comparisons. This gives us four groups for comparison: sham male, concussed male, sham female, and concussed female. A major point to note with this level of statistical testing is that by increasing the number of groups, the number of comparisons also increases, which decreases the statistical power of post-hoc multiple comparisons testing^44^. This is critical to keep in mind, as it can result in a comparison being reported as negative, even though with fewer comparisons, it may have been positive. This phenomenon can be seen in this study. When comparing concussed males to concussed females (Table 4), IL-18 (Figure 5H) was significantly greater in males than females. However, by increasing the number of comparisons with added groups, the difference in IL-18 between concussed males and concussed females is no longer significant. All other significant pairwise comparisons remained significant, even with less power. This suggests that IL-18 may still be an important biomarker to investigate in mTBI, but is not as strongly affected by sex or injury as other proteins in this panel.

Incorporating all four experimental groups into our statistical testing revealed some interesting baseline differences that may prove to be confounding to our previous results (Table 5). First, the sham male group was shown to have significantly higher levels of CXCL3 (Figure 6A) and VEGF (Figure 6G) at 3 dpi compared to sham female and significantly higher levels of CXCL3 (Figure 6A) and CXCL2 (Figure 6B) at 7 dpi than concussed female. Also the shame male group was shown to have significantly lower levels of IFNγ (Figure 6C) at 7 dpi than sham female and concussed female. These results likely do not affect interpretation of directly comparing concussed males and concussed females (Table 4), but instead affect interpretation of the injury responses separated by sex (Table 2 & 3). For example, sham males exhibited significantly lower levels of IFNγ (Figure 6C) compared to both sham female and concussed female at 7 dpi, but there was no statistical significance between concussed male and sham female, concussed female, or sham male. It’s possible that the lack of significance in this comparison is a consequence of the limited study size, and a larger study would be able to confirm whether or not there is a significant difference in IFNγ.

### 4.6. Presence of microbleeds corresponds with increase in Iba1+ microglia number

Despite a concussion being a mild traumatic brain injury, there are still noticeable differences in brain pathology between sexes after a concussion. We performed histological characterization using a quantitative and semi-quantitative methodology to assess the presence of Iba1+ microglia (Figure 7) and GFAP+ astrocytes (Figure S1), as well as microbleeds (Figure 8), respectively. Stereological quantification of Iba1+ microglia, in the corpus callosum shows a significant increase in the estimated total number of microglia in the corpus callosum of males at 14 dpi compared to 7 dpi (Figure 7G). Stereological quantification of GFAP+ astrocytes in the corpus callosum showed no significant differences between any groups. The overall trend of this data corresponds well with the semi-quantitative scoring of microbleed presence, where each instance of a microbleed is graded on a scale of 0-3 (0 = none, 1 = a few granules, 2 = several granules, 3 = a lot of granules). This scoring reveals that males experience a significantly greater presence of microbleeds at 14 dpi compared to male sham, males at 7 dpi, and females at 14 dpi (Figure 8G). This correlation makes sense, as microglia are known to phagocytose blood products after hemorrhage^45,46^. To confirm this response, co-staining of Iba1 and Prussian Blue was performed and showed multiple Iba1+ microglia phagocytosed microbleed byproducts (Figure 9). This data suggests the possibility that the pro-inflammatory CSF protein expression seen in males after a concussion could be driven by an increase in pro-inflammatory microglia polarization due to a increase in BBB permeability. Future studies should investigate variability in microglial phagocytosis between sexes as a possible mechanism underlying the sex-dependent differences in neuroinflammation after concussion.

### 4.7. Limitations

This study was designed with three goals. First, to establish a method of minimally invasive serial CSF collection that could be used by any laboratory without specialized tools to enable future longitudinal studies utilizing clinically relevant outputs. Second, to determine if the serial collection of CSF altered the CSF proteome in the absence of an experimental condition. Third, to establish a framework for interrogating sex differences in levels of biomarkers for mTBI in rats. While this study accomplishes all three goals, it is not without its limitations. First, the sample size in the present study is small. Each group at each timepoint has n=4-6 rats. While group numbers are sufficient for statistical analysis, groups of 10-12 rats would be ideal and would allow for testing of behavioral outcomes in future interventional studies. Second, it is extremely likely that there are other proteins that play a significant role in mTBI inflammation that are not addressed in this study. This is mainly due to the lack of analytes commercially available in multiplex immunoassay kits for rats. As more and more kits designed for rats come to market, studies like this will be able to be expanded. Last, a limitation of this study is the absence of behavioral data. Future studies should aim to correlate clinically relevant biomarkers over time with functional outputs like behavior to maximize the amount of data that can be generated with the least number of animals.

## 5. Conclusion

This study was designed to accomplish three goals. First, to describe a method of minimally invasive serial CSF collection. The method described in this study can easily be adapted by any laboratory prepared for animal studies. There is no need for special tools or custom-built equipment. Second, to confirm that serial collection of the CSF does not alter CSF protein levels. Of the 17 analytes we tested, 16 showed no difference between serial collection and single collection. TIMP-1 was significantly increased which is expected as TIMP-1 is involved in alleviating inflammatory pain related to wound healing. Third, to establish a framework for assessing sex differences in neuroinflammation after a concussion. We showed that results vary based on the framing of the statistical test. However, it is evident that males experience a more robust inflammatory response to a single concussion than females, possibly caused by increased BBB permeability and microglial polarization. Future studies should adopt this method of serial CSF collection to perform longitudinal studies to better understand the temporal dynamics of neuroinflammation not just after neurotrauma, but any neurological disorder.

## Acknowledgements

We want to thank technical staff at the Sue & Bill Gross Stem Cell Center, especially Rebecca Nishi, for her help with animal studies. We would like to thank Dr. Dequina Nicholas and the Nicholas lab for their generous support with all Luminex assays.

## Ethics Declaration

All experiments, including animal housing conditions, surgical procedures, and postoperative care, were in accordance with the Institutional Animal Care and Use Committee guidelines at the University of California–Irvine (UCI).

## Data Availability Statement

The datasets generated during and/or analysed during the current study are available from the corresponding author on reasonable request.

---

**Figure S1:**
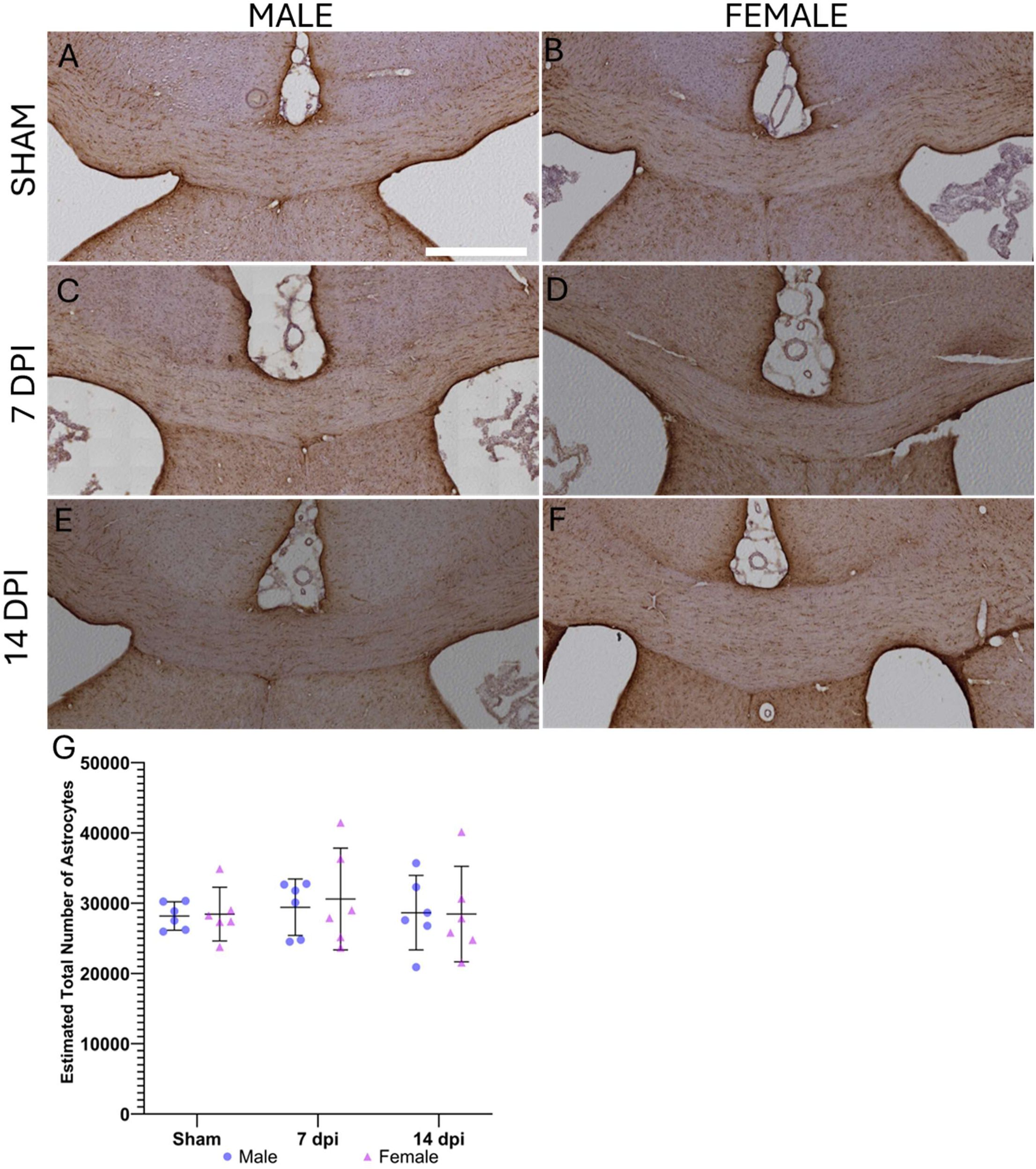
There are no differences in the number of astrocytes across groups. Stereological quantification of GFAP+ (brown) cells counterstained with hematoxylin (blue) was performed on serial sections of brain tissue. Representative images are shown (A-F). Quantification was performed only in the corpus callosum and is reported as the estimated total number of astrocytes (G). Data is presented as mean with 95% Cl. Statistical analysis was performed using a two-way ANOVA and post-hoc Holm-Sidak multiple comparisons test. Scale bar = 100 μm.

**Figure S2:**
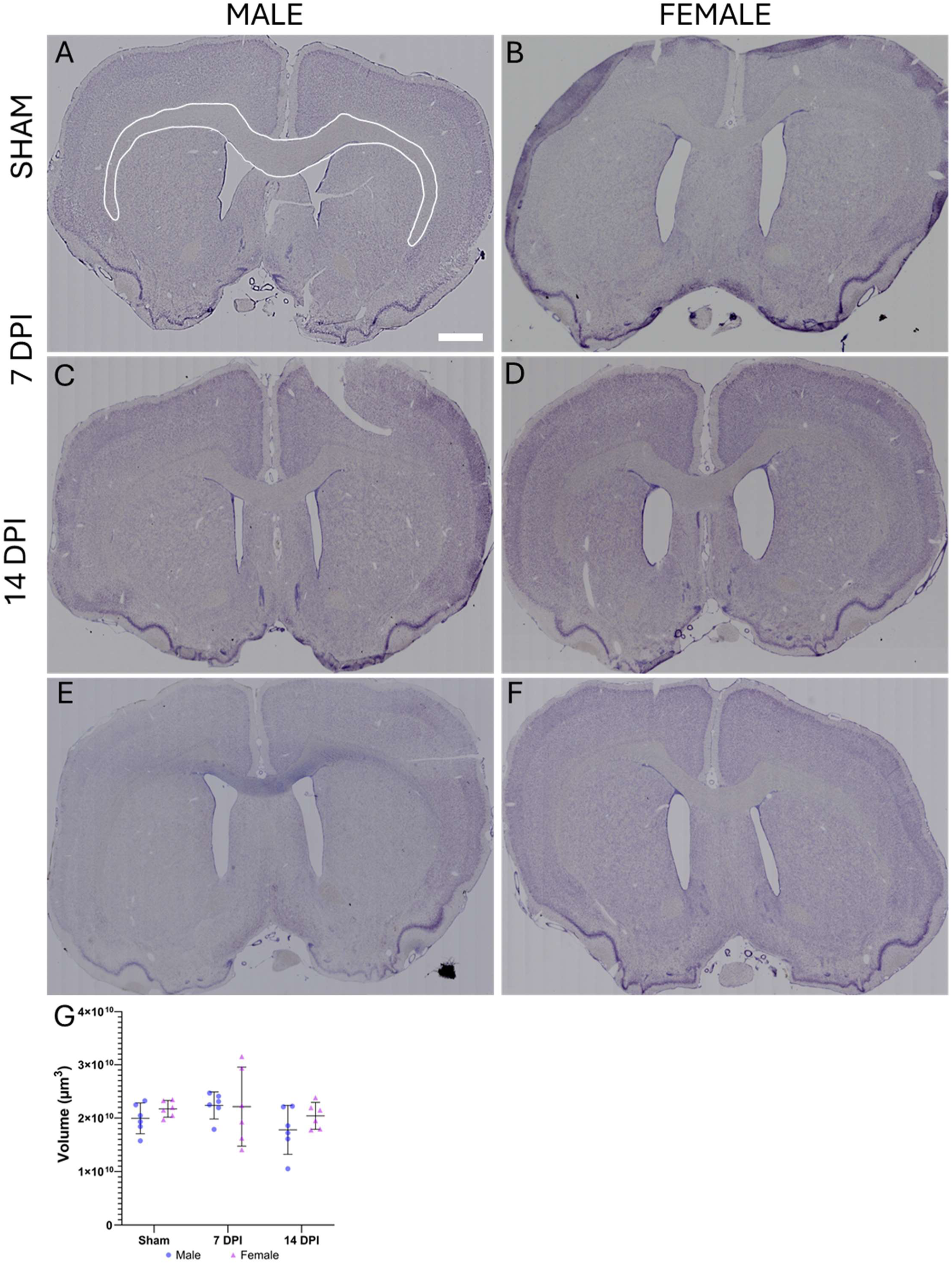
There are no differences in corpus callosum volume across groups. Cavalieri estimation was performed on serial sections of brain tissue to estimate volume of corpus callosum. Representative images are shown (A-F). An example ROI for corpus callosum volume estimation can be seen in A, depicted with a white outline. Volume estimations are presented as mean with 95% Cl (G). Statistical analysis was performed using a two-way ANOVA and post-hoc Holm-Sidak multiple comparisons test. Scale bar = 1000 μm.

## References

1 Guan B, Anderson DB, Chen L, Feng S, Zhou H. Global, regional and national burden of traumatic brain injury and spinal cord injury, 1990-2019: a systematic analysis for the Global Burden of Disease Study 2019. BMJ Open 2023;13:. 10.1136/BMJOPEN-2023-075049.

2 Capizzi A, Woo J, Verduzco-Gutierrez M. Traumatic Brain Injury An Overview of Epidemiology, Pathophysiology, and Medical Management. Med Clin N Am 2020;104:213–38. 10.1016/j.mcna.2019.11.001.

3 Levy AM, Saling MM, Anderson JFI. Frequency and extent of cognitive complaint following adult civilian mild traumatic brain injury: A systematic review and meta-analysis. Brain Impair 2023:309–32. 10.1017/BrImp.2022.19.

4 Grovola MR, Von Reyn C, Loane DJ, Cullen DK. Understanding microglial responses in large animal models of traumatic brain injury: an underutilized resource for preclinical and translational research. J Neuroinflammation 2023;20:67. 10.1186/s12974-023-02730-z.

5 Neale KJ, Reid HMO, Sousa B, McDonagh E, Morrison J, Shultz S, et al. Repeated mild traumatic brain injury causes sex-specific increases in cell proliferation and inflammation in juvenile rats. J Neuroinflammation 2023;20:. 10.1186/S12974-023-02916-5.

6 Fraunberger EA, Dejesus P, Zanier ER, Shutt TE, Esser MJ. Acute and Persistent Alterations of Cerebellar Inflammatory Networks and Glial Activation in a Rat Model of Pediatric Mild Traumatic Brain Injury. J Neurotrauma 2020;37:1315–30. 10.1089/neu.2019.6714.

7 Feigin VL, Nichols E, Alam T, Bannick MS, Beghi E, Blake N, et al. Global, regional, and national burden of neurological disorders, 1990–2016: a systematic analysis for the Global Burden of Disease Study 2016. Lancet Neurol 2019;18:459–80.

8 Gold EM, Vasilevko V, Hasselmann J, Tiefenthaler C, Hoa D, Ranawaka K, et al. Repeated Mild Closed Head Injuries Induce Long-Term White Matter Pathology and Neuronal Loss That Are Correlated With Behavioral Deficits. ASN Neuro 2018;10:. 10.1177/1759091418781921/ASSET/IMAGES/LARGE/10.1177_1759091418781921-FIG2.JPEG.

9 Gard A, Vedung F, Piehl F, Khademi M, Wernersson MP, Rorsman I, et al. Cerebrospinal fluid levels of neuroinflammatory biomarkers are increased in athletes with persistent post-concussive symptoms following sports-related concussion. J Neuroinflammation 2023;20:1–10. 10.1186/S12974-023-02864-0/FIGURES/2.

10 Visser K, de Koning ME, Ciubotariu D, Kok MGJ, Sibeijn-Kuiper AJ, Bourgonje AR, et al. An exploratory study on the association between blood-based biomarkers and subacute neurometabolic changes following mild traumatic brain injury. J Neurol 2023. 10.1007/s00415-023-12146-7.

11 Giza CC, Hovda DA. The new neurometabolic cascade of concussion. Neurosurgery 2014;75:S24–33. 10.1227/NEU.0000000000000505.

12 Howell DR, Southard J. The Molecular Pathophysiology of Concussion. Clin Sports Med 2021;40:39–51. 10.1016/J.CSM.2020.08.001.

13 Shultz SR, Mcdonald SJ, Haar CV, Meconi A, Vink R, Van Donkelaar P, et al. The potential for animal models to provide insight into mild traumatic brain injury: Translational challenges and strategies. Neurosci Biobehav Rev 2017;76:396–414. 10.1016/j.neubiorev.2016.09.014.

14 Ryan E, Kelly L, Stacey C, Huggard D, Duff E, McCollum D, et al. Mild-to-severe traumatic brain injury in children: altered cytokines reflect severity. J Neuroinflammation 2022;19:36. 10.1186/s12974-022-02390-5.

15 Villapol S, Loane DJ, Burns MP. Sexual dimorphism in the inflammatory response to traumatic brain injury. Glia 2017;65:1423–38. 10.1002/glia.23171.

16 Daisy CC, Varinos S, Howell DR, Kaplan K, Mannix R, Meehan WP, et al. Proteomic Discovery of Noninvasive Biomarkers Associated With Sport-Related Concussions. Neurology 2022;98:E186–98. 10.1212/WNL.0000000000013001.

17 Brotfain E, Gruenbaum SE, Boyko M, Kutz R, Zlotnik A, Klein M. Neuroprotection by Estrogen and Progesterone in Traumatic Brain Injury and Spinal Cord Injury. Curr Neuropharmacol 2016;14:641. 10.2174/1570159X14666160309123554.

18 Liu S (Steve), Pickens S, Barta Z, Rice M, Dagher M, Lebens R, et al. Neuroinflammation drives sex-dependent effects on pain and negative affect in a murine model of repeated mild traumatic brain injury. Pain 2023. 10.1097/j.pain.0000000000003084.

19 Beckmann L, Obst S, Labusek N, Abberger H, Köster C, Klein-Hitpass L, et al. Regulatory T Cells Contribute to Sexual Dimorphism in Neonatal Hypoxic-Ischemic Brain Injury. Stroke 2022;53:381–90. 10.1161/STROKEAHA.121.037537/FORMAT/EPUB.

20 Chakrabarti M, Das A, Samantaray S, Smith JA, Banik NL, Haque A, et al. Molecular mechanisms of estrogen for neuroprotection in spinal cord injury and traumatic brain injury. Rev Neurosci 2016;27:271–81. 10.1515/revneuro-2015-0032.

21 Wang J, Hou Y, Zhang L, Liu M, Zhao J, Zhang Z, et al. Estrogen Attenuates Traumatic Brain Injury by Inhibiting the Activation of Microglia and Astrocyte-Mediated Neuroinflammatory Responses. Mol Neurobiol 2021;58:1052–61. 10.1007/s12035-020-02171-2.

22 Zlotnik A, Leibowitz A, Gurevich B, Ohayon S, Boyko M, Klein M, et al. Effect of estrogens on blood glutamate levels in relation to neurological outcome after TBI in male rats. Intensive Care Med 2012;38:137–44. 10.1007/S00134-011-2401-3.

23 Wang KK, Yang Z, Zhu T, Shi Y, Rubenstein R, Tyndall JA, et al. An update on diagnostic and prognostic biomarkers for traumatic brain injury. Expert Rev Mol Diagn 2018:165–80. 10.1080/14737159.2018.1428089.

24 Liu MC, Akinyi L, Scharf D, Mo J, Larner SF, Muller U, et al. Ubiquitin C-terminal hydrolase-L1 as a biomarker for ischemic and traumatic brain injury in rats. Eur J Neurosci 2010;31:722–32. 10.1111/j.1460-9568.2010.07097.x.

25 Yao C, Williams AJ, Ottens AK, Lu XCM, Liu MC, Hayes RL, et al. P43/pro-EMAPII: A potential biomarker for discriminating traumatic versus ischemic brain injury. J Neurotrauma 2009;26:1295–305. 10.1089/neu.2008.0811.

26 Zoltewicz JS, Mondello S, Yang B, Newsom KJ, Kobeissy F, Yao C, et al. Biomarkers track damage after graded injury severity in a rat model of penetrating brain injury. J Neurotrauma 2013;30:1161–9. 10.1089/NEU.2012.2762/ASSET/IMAGES/LARGE/FIGURE5.JPEG.

27 Han JR, Yang Y, Wu TW, Shi TT, Li W, Zou Y. A Minimally-Invasive Method for Serial Cerebrospinal Fluid Collection and Injection in Rodents with High Survival Rates. Biomedicines 2023;11:. 10.3390/biomedicines11061609.

28 Baskin BM, Logsdon AF, Janet Lee S, Foresi BD, Peskind E, Banks WA, et al. Timing matters: Sex differences in inflammatory and behavioral outcomes following repetitive blast mild traumatic brain injury. Brain Behav Immun 2023;110:222–36. 10.1016/j.bbi.2023.03.003.

29 Heyburn L, Batuure A, Wilder D, Long J, Sajja VS. Neuroinflammation Profiling of Brain Cytokines Following Repeated Blast Exposure. Int J Mol Sci 2023, Vol 24, Page 12564 2023;24:12564. 10.3390/IJMS241612564.

30 Breen EJ, Tan W, Khan A. The Statistical Value of Raw Fluorescence Signal in Luminex xMAP Based Multiplex Immunoassays. Sci Rep 2016;6:. 10.1038/SREP26996.

31 Breen EJ, Polaskova V, Khan A. Bead-based multiplex immuno-assays for cytokines, chemokines, growth factors and other analytes: median fluorescence intensities versus their derived absolute concentration values for statistical analysis. Cytokine 2015;71:188–98. 10.1016/J.CYTO.2014.10.030.

32 Batllori M, Casado M, Sierra C, Salgado M del C, Marti-Sanchez L, Maynou J, et al. Effect of blood contamination of cerebrospinal fluid on amino acids, biogenic amines, pterins and vitamins. Fluids Barriers CNS 2019;16:1–9. 10.1186/S12987-019-0154-5/TABLES/2.

33 Knight BE, Kozlowski N, Havelin J, King T, Crocker SJ, Young EE, et al. TIMP-1 Attenuates the Development of Inflammatory Pain Through MMP-Dependent and Receptor-Mediated Cell Signaling Mechanisms. Front Mol Neurosci 2019;12:. 10.3389/FNMOL.2019.00220/FULL.

34 Giza CC, McCrea M, Huber D, Cameron KL, Houston MN, Jackson JC, et al. Assessment of Blood Biomarker Profile After Acute Concussion During Combative Training Among US Military Cadets: A Prospective Study From the NCAA and US Department of Defense CARE Consortium. JAMA Netw Open 2021;4:E2037731. 10.1001/JAMANETWORKOPEN.2020.37731.

35 Hicks SD, Onks C, Kim RY, Zhen KJ, Loeffert J, Loeffert AC, et al. Refinement of saliva microRNA biomarkers for sports-related concussion. J Sport Heal Sci 2023;12:369–78. 10.1016/j.jshs.2021.08.003.

36 Vedantam A, Brennan J, Levin HS, McCarthy JJ, Dash PK, Redell JB, et al. Early versus Late Profiles of Inflammatory Cytokines after Mild Traumatic Brain Injury and Their Association with Neuropsychological Outcomes. J Neurotrauma 2021;38:53–62. 10.1089/neu.2019.6979.

37 Amen EM, Brecheisen M, Sach-Peltason L, Bergadano A. Refinement of a model of repeated cerebrospinal fluid collection in conscious rats. Lab Anim 2017;51:44–53. 10.1177/0023677216646069/ASSET/IMAGES/LARGE/10.1177_0023677216646069-FIG2.JPEG.

38 Chaban V, Clarke GJB, Skandsen T, Islam R, Einarsen CE, Vik A, et al. Systemic Inflammation Persists the First Year after Mild Traumatic Brain Injury: Results from the Prospective Trondheim Mild Traumatic Brain Injury Study. J Neurotrauma 2020;37:2120–30. 10.1089/neu.2019.6963.

39 Barreto GE, Thelin EP, Zhang Y, Madathil SK, Wilfred BS, Urankar SE, et al. Early Microglial Activation Following Closed-Head Concussive Injury Is Dominated by Pro-Inflammatory M-1 Type. Front Neurol 2018;9:414936. 10.3389/FNEUR.2018.00964.

40 Simon DW, McGeachy MJ, Baylr H, Clark RSB, Loane DJ, Kochanek PM. The far-reaching scope of neuroinflammation after traumatic brain injury. Nat Rev Neurol 2017:171–91. 10.1038/nrneurol.2017.13.

41 Miao W, Zhao Y, Huang Y, Chen D, Luo C, Su W, et al. IL-13 Ameliorates Neuroinflammation and Promotes Functional Recovery after Traumatic Brain Injury. J Immunol 2020;204:1486–98. 10.4049/jimmunol.1900909.

42 Lladó J, Tolosa L, Olmos G. Cellular and molecular mechanisms involved in the neuroprotective effects of VEGF on motoneurons. Front Cell Neurosci 2013;7:64456. 10.3389/FNCEL.2013.00181/BIBTEX.

43 Di Battista AP, Churchill N, Rhind SG, Richards D, Hutchison MG. The relationship between symptom burden and systemic inflammation differs between male and female athletes following concussion. BMC Immunol 2020;21:. 10.1186/s12865-020-0339-3.

44 Lee S, Lee DK. What is the proper way to apply the multiple comparison test? Korean J Anesthesiol 2018;71:353–60. 10.4097/kja.d.18.00242.

45 Yu F, Wang Y, Stetler AR, Leak RK, Hu X, Chen J. Phagocytic microglia and macrophages in brain injury and repair. CNS Neurosci Ther 2022;28:1279–93. 10.1111/cns.13899.

46 Chen W, Zhang Y, Zhai X, Xie L, Guo Y, Chen C, et al. Microglial phagocytosis and regulatory mechanisms after stroke. J Cereb Blood Flow Metab 2022;42:1579–96. 10.1177/0271678X221098841.

